# Integrative host transcriptomic and mucosal microbiome profiling reveals region-specific host-microbiome associations across the human intestine

**DOI:** 10.64898/2026.05.13.725025

**Authors:** Erica P. Ryu, Cheryl A. Keller, Robert G. Nichols, Hanh N. Tran, Patricia R. Brocious, Leonard R. Harris, Walter A. Koltun, Gregory S. Yochum, Emily R. Davenport

**Affiliations:** Department of Biology, Pennsylvania State University, University Park, PA; Department of Biochemistry and Molecular Biology, University Park, PA; Genomics Research Incubator, University Park, PA; Huck Institutes of the Life Sciences, University Park, PA; Department of Surgery, Division of Colon & Rectal Surgery, Pennsylvania State University College of Medicine, Hershey, PA; Department of Molecular and Precision Medicine, Pennsylvania State University College of Medicine, Hershey, PA

## Abstract

Host genetics shapes gut microbiome composition, yet the physiological mechanisms underlying this relationship remain poorly understood. Characterizing associations between host gene expression and the mucosal microbiome offers a promising route to identifying the host pathways and microbial taxa most likely to interact physiologically. However, existing investigations have been conducted primarily in acute disease contexts and within the colon, leaving host-microbiome associations outside of acute inflammatory contexts and those in undersampled regions such as the terminal ileum poorly characterized. To address these gaps, we profiled paired host gene expression from full-thickness resections and mucosal microbiome data, both from macroscopically non-inflamed tissue from Crohn’s disease patients undergoing surgery across three intestinal sites: terminal ileum (n = 32), cecum (n = 35), and right colon (n = 30). Using a multi-level analytical framework including Procrustes analysis, sparse canonical correlation analysis, and elastic net regression, we identified significant associations between the mucosal transcriptome and microbiome. Intestine-wide, genes enriched in immune and intestinal barrier integrity pathways were associated with heritable taxa including *Fusicatenibacter*, consistent with patterns observed in microbiome genome-wide association studies. Region-specific analysis identified the terminal ileum as a distinct site of host-microbiome interaction, with associations involving metabolic and barrier-related pathways not observed in the large intestine. Notable terminal ileum-specific associations included *PCDH20* with *Faecalitalea* and *ACAT1* with *Lactococcus*, implicating epithelial barrier maintenance and host-microbiome metabolic interactions, respectively. These findings advance our understanding of the physiological basis of host-microbiome interactions across the intestine.

**Importance:** The human gut is home to trillions of microorganisms that interact with the intestinal lining, yet we have a limited understanding of the specific biological processes involved in these interactions. Most studies characterizing the relationships between host gene expression and the gut microbiome have focused on the colon and on active disease contexts, leaving it unclear whether the associations observed reflect fundamental host-microbiome biology or disease-specific responses. By examining mucosal tissue, where host cells and microbes are in direct contact, across three sites in non-acutely inflamed tissue, we show that expression of immune defense and barrier maintenance genes is broadly associated with the microbiome across the intestine. We also identify distinct classes of associations in the terminal ileum, including host genes involved in metabolic processes. These findings provide a foundation for understanding how host biology and the gut microbiome are linked outside of acute disease.

## Introduction

The gut microbiome exists in constant dialogue with its host, and understanding the physiological basis of this relationship remains a fundamental challenge in biology and medicine. Host genetics is a well-established driver of gut microbiome composition, yet the mechanisms by which this occurs remain unclear (Bonder et al., 2016; Davenport et al., 2015; Goodrich et al., 2016; Lim et al., 2017; Lopera-Maya et al., 2022; Qin et al., 2022; Turpin et al., 2016; J. Wang et al., 2016). Several taxa, such as *Akkermansia,* Christensenellaceae, and *Bifidobacterium*, are particularly notable for their pattern of heritability and associations with disease outcomes (Ghaffari et al., 2022; Hidalgo-Cantabrana et al., 2017), yet we still lack an understanding of what host processes underlie this heritability. Host gene expression represents a dynamic intermediate between the genome and the microbial environment, shaped by both genetic variation and environmental signals. Characterizing host transcriptome-microbiome associations therefore offers a promising route to uncover the physiological mechanisms governing this bidirectional relationship: how the host shapes its microbial community and how the microbiome in turn influences host gene expression (Bubier et al., 2021; Ferretti et al., 2025; Nichols & Davenport, 2021). Yet despite this potential, the physiological basis of host transcriptome-microbiome associations remains poorly characterized outside of disease contexts and across intestinal regions.

Integrating host gene expression and microbiome data enables hypothesis-free, large-scale identification of associations across these two data modalities, offering biological insights into host-microbiome interactions. Indeed, several studies have applied this approach and identified consistent associations between the host transcriptome and microbiome. Host immune genes and pathways have consistently emerged in host-microbiome associations across multiple investigations (Cai et al., 2021; Hu et al., 2024; Lloyd-Price et al., 2019; Priya et al., 2022).

However, these studies have been conducted largely in the context of inflammatory bowel disease (IBD), irritable bowel syndrome (IBS), and colorectal cancer, either between healthy controls and samples with disease or between non-inflamed and inflamed tissue samples. While it is certainly possible that intestinal genes generally associate with microbes outside of disease, it is difficult to fully disentangle host-microbiome associations from the effect of disease. For example, in a study involving colorectal cancer, the genes primarily found to be associated with the microbes were tumor suppressors (Kim et al., 2022), suggesting context-dependent associations that make it challenging to discern what associations exist between the host transcriptome and microbiome outside of acute disease. A study conducted in non-acutely inflamed tissue is therefore needed to characterize baseline host-microbiome associations.

A further limitation of existing work is its reliance on sample types that do not fully capture relevant biology. Fecal samples have been commonly used as the proxy for the gut microbiota and blood for gene expression to identify associations between immune genes, such as IL-1β and IL-2 and gut microbes; however, these sample types offer limited resolution for identifying the specific genes and microbes interacting within the gut (Jangi et al., 2016; Zhou et al., 2019). While convenient, the fecal microbiome is primarily composed of luminal microbes, which differ substantially from the mucosal microbiome (Carroll et al., 2010; Chen et al., 2012; Rangel et al., 2015). It has been hypothesized that the mucosal microbiome may show stronger associations with host genetics and disease processes (Juge, 2022). Tissue biopsies and surgical resections offer a more direct route to characterizing the mucosal microbiome (Q. Tang et al., 2020), though the challenges of cleanly acquiring these sample types have limited their use and left mucosal microbiome interactions relatively underexplored.

Additionally, most host transcriptome-microbiome studies have focused on the colon or sites more easily accessible by endoscopy, leaving higher regions of the gastrointestinal tract largely undercharacterized. Regions along the intestine function differently and house distinct microbial communities (Burclaff et al., 2022; Chavez-Arroyo et al., 2025; Hickey et al., 2023; Yang et al., 2025). The small intestine is typically lower in microbial abundance and diversity compared to the large intestine, likely due to differences in transit time (3-5 hours in the small bowel vs >30 hours in the colon) (Martinez-Guryn et al., 2019), though diversity increases toward the distal end of the small bowel (Kastl et al., 2019). While the terminal ileum is suspected to be an area of high microbial activity, there is no consensus on its microbial composition. Some studies report it as distinct from the colon, whereas others do not (Dave et al., 2011; Villmones et al., 2018; M. Wang et al., 2005). Moreover, it is unclear whether trends observed in the luminal microbiome hold for the mucosal microbiome (Li et al., 2015). Acquiring samples from these sites requires surgical resection, limiting most prior investigations to the colon (Jensen et al., 2023; Leite et al., 2020; Q. Tang et al., 2020). Although a handful of prior studies have included ileal samples, this region was either not investigated in depth or location was regressed out (Hu et al., 2024; Lloyd-Price et al., 2019). As a result, host transcriptome-microbiome associations in the ileum remain undercharacterized, despite the growing interest in the small intestine as a site of gastrointestinal disease and a source of potential therapeutic targets (Ruigrok et al., 2023; Yersin & Vonaesch, 2024).

To address these gaps, we investigated relationships between host gene expression and the mucosal microbiome along the lower gastrointestinal tract, using macroscopically non-inflamed tissue to minimize the confounding effects of active disease and gain insights into fundamental physiological host-microbiome relationships. We examined paired host RNA-seq and mucosal microbiome data from three intestinal sites – terminal ileum, cecum, and right colon – that are underrepresented in the literature due to their difficulty in being obtained. Using a multi-level analytical framework, we demonstrate that host genes enriched in immune pathways and intestinal barrier function are associated with heritable taxa, providing transcriptomic insight into the mechanisms that may underlie host genetic influence on the microbiome. Furthermore, we show that the terminal ileum is an area of distinct host gene expression-microbe activity, with site-specific associations involving genes enriched in pathways for metabolism and intestinal barrier function, suggesting this undersampled site may be an important locus of host-microbiome interaction.

## Results

### Description of cohort

We examined the microbiomes and transcriptomes of human tissue samples originating from three different intestinal locations: terminal ileum (n = 50 microbiome, n = 39 RNA-seq, n = 32 overlapping), cecum (n = 43 microbiome, n = 44 RNA-seq, n = 35 overlapping), and right colon (n = 39 microbiome, n = 37 RNA-seq, n = 30 overlapping, **Figure 1**). We selected these three sites as they have been underrepresented in the literature, likely due to the difficulty in obtaining them, as well as their differing physiological functions. Because these intestinal sites are difficult to access via standard colonoscopy, we used tissue collection acquired through surgical resection. Samples were therefore obtained from Crohn’s disease patients undergoing colectomy, a design that provided access to otherwise unobtainable tissue while leveraging an existing surgical cohort. Critically, to minimize the confounding effects of active inflammation and better approximate a non-acute disease state, only macroscopically non-inflamed tissue was selected for analysis. Samples from sites of acute IBD flares were excluded.

**Figure 1:**
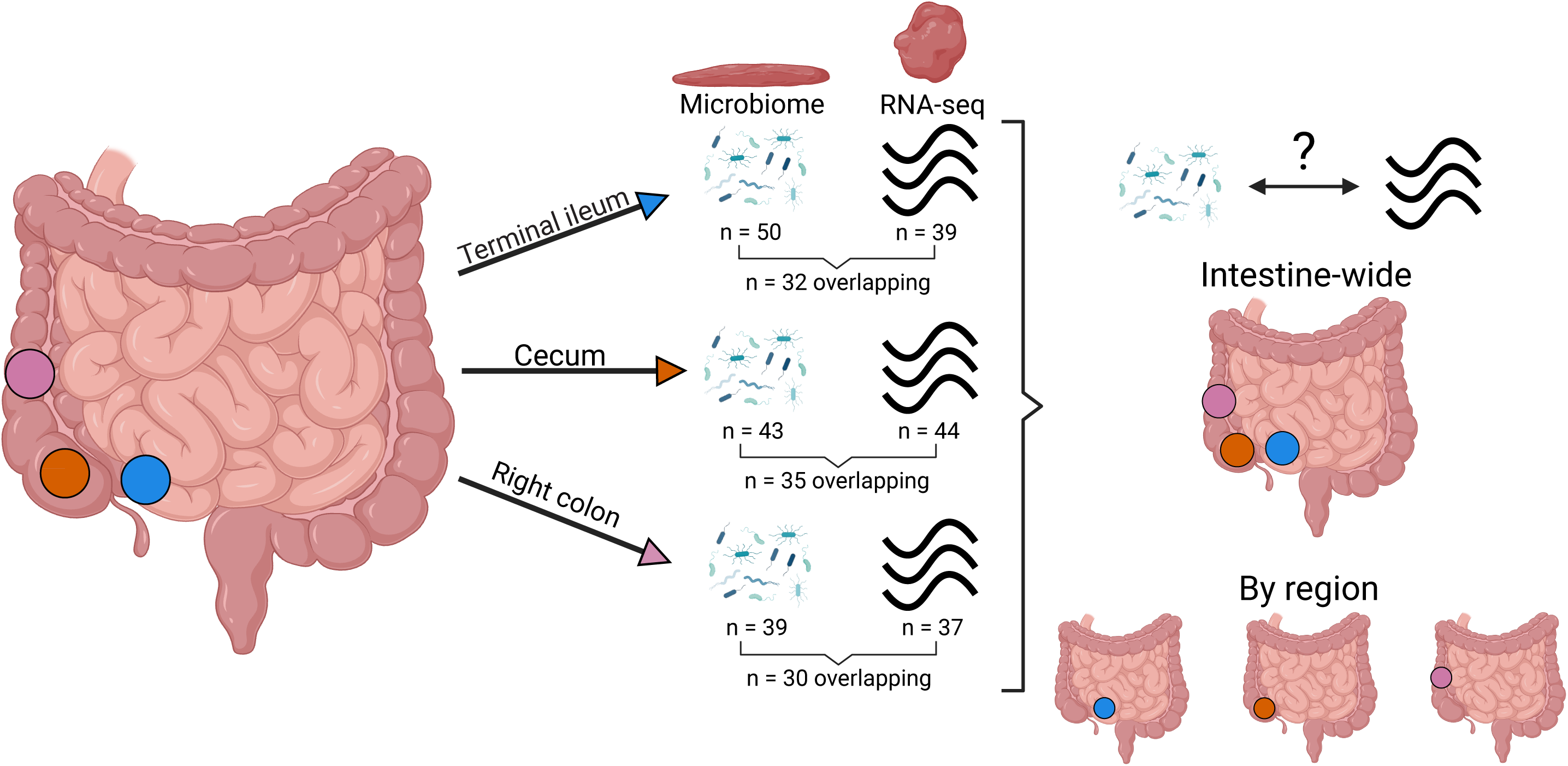
Intestinal regions sampled and study workflow. Overview of the study design. Paired macroscopically non-inflamed full-thickness tissue and mucosal scrapings were obtained from the terminal ileum (blue), cecum (orange), and right colon (pink) from Crohn’s disease patients. 16S rRNA gene microbiome data were generated from mucosal scrapings and host RNA-seq data were generated from full-thickness tissue. These data were used to identify gene expression-microbiome associations both across the intestine and within individual intestinal regions.

Although the samples were classified as macroscopically non-inflamed tissue, they originated from patients with Crohn’s disease. While Crohn’s disease is characterized by localized acute flares, there can be subclinical systemic inflammation in non-inflamed regions that could affect microbiome or transcriptional profiles. To evaluate whether such effects were detectable, non-cancerous tissue samples from the right colon collected from individuals with colorectal cancer were included as a comparison (n = 18 microbiome, n = 15 RNA-seq, n = 14 overlapping), as colorectal cancer generally does not cause underlying systemic inflammation akin to Crohn’s disease (Bhat et al., 2022). Microbiome comparisons revealed no significant differences in diversity (p > 0.05, ANOVA; **Supplemental Figure 1A**) or composition (p = 0.28, PERMANOVA; **Supplemental Figure 1B**) between groups. Similarly, no differentially expressed genes were identified between colorectal cancer and Crohn’s disease patients (**Supplemental Figure 2**). These results suggest that large-scale transcriptional or compositional differences attributable to underlying inflammation are unlikely in our macroscopically non-inflamed Crohn’s disease samples, though we cannot exclude the possibility of subtler effects. These samples therefore serve as a reasonable approximation of a non-acute disease context for the purposes of this study. Subsequent analyses solely involved the use of samples from Crohn’s disease patients.

The microbiome differs between intestinal locations in both mouse models and humans (Lkhagva et al., 2021; Shalon et al., 2023). However, there is concern about cross-contamination between regions, as human samples have typically been acquired via colonoscopy or endoscopy (Q. Tang et al., 2020). To address this limitation, we used surgical tissue resections and matched mucosal scrapings to minimize cross-contamination arising from sample acquisition. Furthermore, as surgical resections are low biomass microbiome samples, extensive measures were implemented to reduce environmental, host, and cross-contamination of the microbiome (see Methods). This sampling scheme allowed us to compare the microbiome between locations to provide a baseline profile of microbiome diversity and composition while minimizing cross-site and environmental contamination concerns.

### Microbiome composition and gene expression differ across intestinal locations

We first characterized microbiome diversity and composition across intestinal locations. We observed no significant differences in mucosal microbiome diversity across locations (p > 0.05, ANOVA; **Figure 2A**), though microbiome composition differed significantly by location (p = 0.0027, PERMANOVA; **Figure 2B**). These differences were primarily driven by the terminal ileum being distinct from the large bowel regions (terminal ileum vs. cecum: p = 0.01, terminal ileum vs. right colon: p = 0.002; pairwise PERMANOVA).

**Figure 2:**
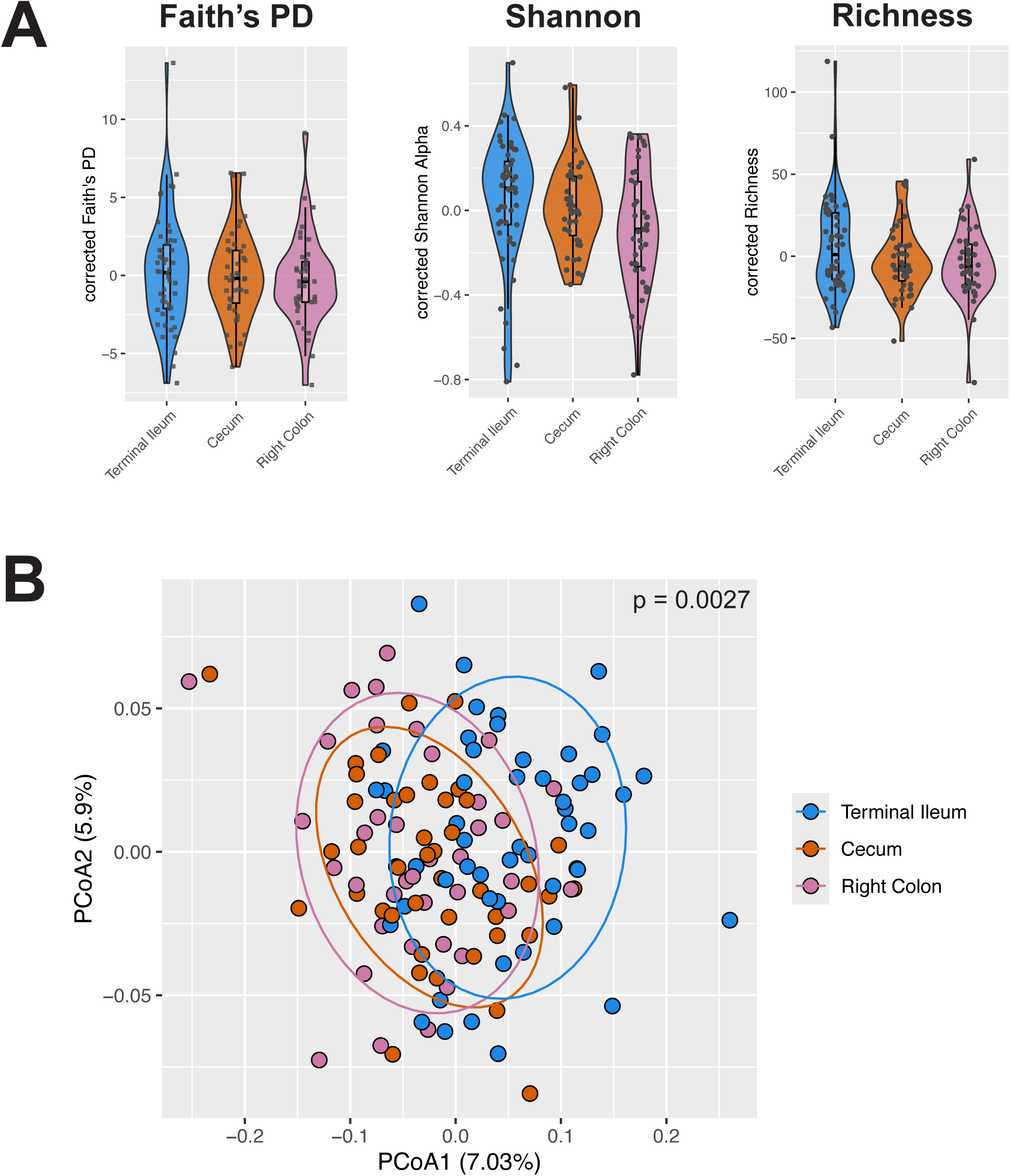
Microbiome composition, but not diversity, differs across intestinal locations. A) Covariate-corrected Faith’s phylogenetic diversity (Faith’s PD, *left*), Shannon alpha diversity (*middle*), and richness (*right*) across all individuals, grouped by intestinal location from proximal to distal. Significant differences are not detected across locations for Faith’s PD, Shannon alpha diversity, and richness (p > 0.05 for all, ANOVA). B) Microbiome composition varies significantly with intestinal location (p = 0.0027, PERMANOVA), with terminal ileum differing from the large bowel regions (terminal ileum vs. cecum: p = 0.01, terminal ileum vs. right colon: p = 0.002; pairwise PERMANOVA). The PCoA plot shows individuals projected based on Bray–Curtis distance and colored by location. Bray–Curtis distances were calculated after correcting for the covariates age, sex, extraction batch, library concentration, and patient.

Differential abundance analysis identified only two taxa as significantly differentially abundant across regions: unclassified *Lactobacillus* and *Granulicatella* (q < 0.05, generalized linear mixed model; **Supplemental Table 1**). However, an additional 55 microbes were nominally significant (nominal p < 0.05, generalized linear mixed model; **Supplemental Table 1**), suggesting that the regional compositional differences observed in the PERMANOVA may reflect small shifts in microbial relative abundance across the community that are below our current power to detect after multiple test correction.

We next examined gene expression across intestinal location. Intestinal physiology differs by region and these differences have been resolved to the cell-type and molecular levels (Burclaff et al., 2022; Chavez-Arroyo et al., 2025; Hickey et al., 2023; Yang et al., 2025). While the literature has extensively documented the differences between intestinal regions (M. D. Bates et al., 2002; Comelli et al., 2009; Glebov et al., 2003), we sought to confirm these differences in our cohort for a baseline profile of gene expression. Principal components analysis revealed that gene expression of the terminal ileum was distinctly different from that of the cecum and right colon (**Figure 3A**). This finding was supported by differential expression analysis, in which we observed that 1,318 genes were differentially expressed between the terminal ileum and the cecum and 1,358 genes between the terminal ileum and right colon (adjusted p < 0.05, DESeq2; **Figure 3B, Supplemental Table 2**). In comparison, only 6 genes were differentially expressed between the cecum and the right colon, which was not unexpected given their proximity.

**Figure 3:**
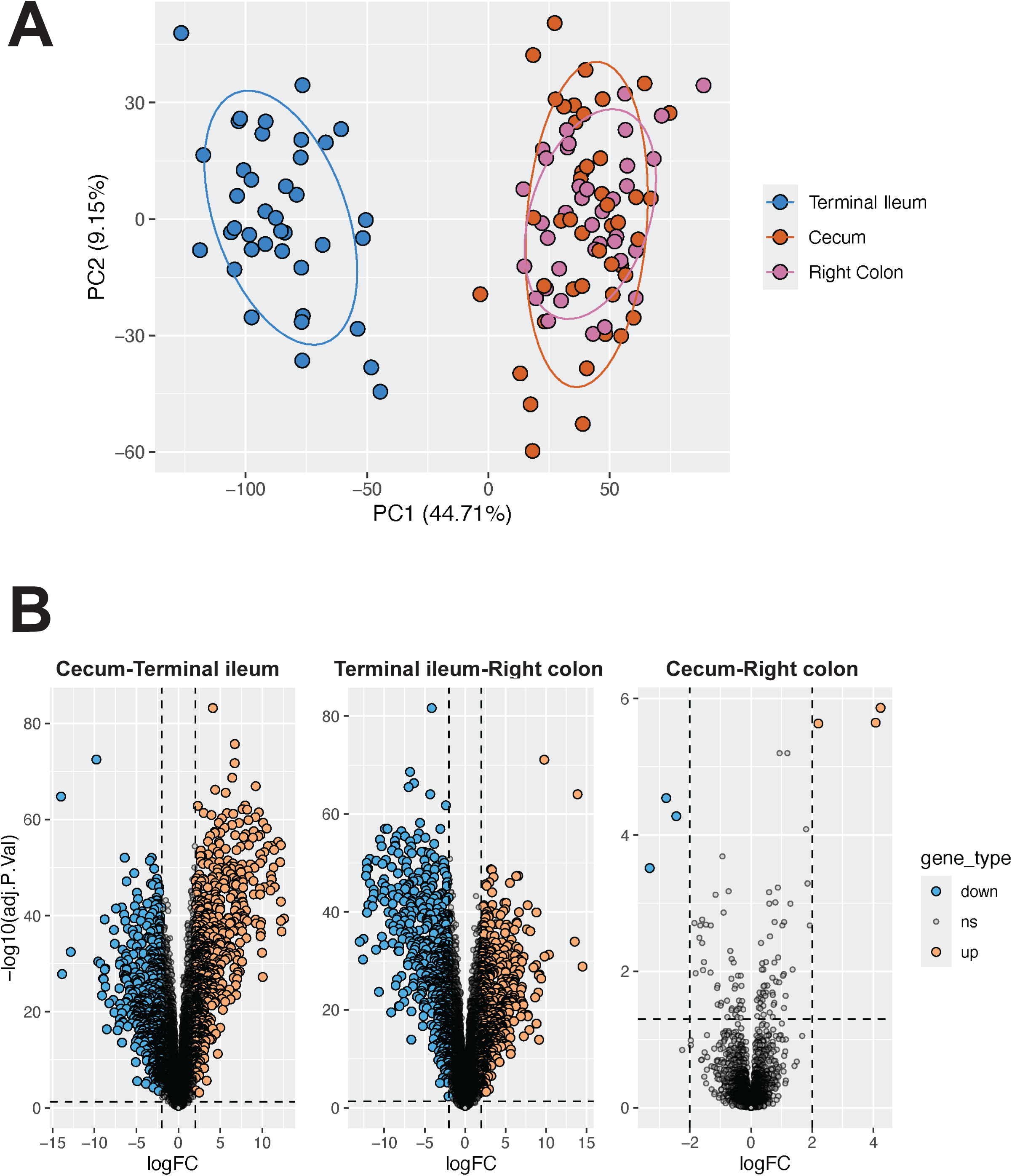
Gene expression differs based on intestinal location. A) PCA plot shows samples colored by intestinal location, with clear separation between the small and large bowel sites as expected. Expression values were corrected for covariates prior to projection, including sex, age, library concentration, RIN score, sequencing batch, and extractor. B) Substantially more genes were differentially expressed between the terminal ileum and cecum (*left*) and the terminal ileum and the right colon (*middle*) compared to the cecum and the right colon (*right*). Genes with log fold changes (logFC) > 2 and Benjamini-Hochberg adjusted p-values < 0.05 were considered significant. For each plot, genes that were more highly expressed in the location listed first are colored in blue, while genes more highly expressed in the location listed second are in orange. For example, in the left plot, genes that were significantly more expressed in the cecum compared to the terminal ileum are blue, while genes more expressed in the terminal ileum compared to the cecum are in orange.

### Expression of host genes enriched in pathways for intestinal barrier integrity and the immune system is associated with gut microbes

With baseline profiles of regional microbiomes and transcriptomes established, we next sought to identify associations between them. While associations between host gene expression and microbes have been observed, these analyses have been limited to specific intestinal regions and conducted primarily in disease contexts (Cai et al., 2021; Hu et al., 2024; Kim et al., 2022; Priya et al., 2022). Our goal was therefore to identify associations between the microbiome and host gene expression across a broader range of intestinal sites. To begin, we identified whether there were any intestine-wide associations between the microbiome and gene expression across the intestine, considering all sites together.

To this end, we employed a three-pronged approach, similar to Priya et al. (Priya et al., 2022). First, we identified whether there was a signal of global concordance between microbiome and transcriptomic profiles. Second, we identified whether there were significant associations between groups of co-expressed genes and groups of co-abundant microbes, motivated by the observation that genes within pathways are often co-regulated and microbes frequently form co-abundance groups. Finally, we identified specific genes associated with specific microbes. Because relationships may exist at any of these levels, this multilevel approach was key for comprehensive characterization of host gene expression-microbe relationships. For example, while a global relationship may not be observed, co-expressed genes and co-abundant microbes may be associated.

We first examined the global patterns of concordance between mucosal microbiome composition and gene expression profiles using Procrustes analysis (**Figure 4A**). Previous work observed a global association only in the context of colorectal cancer, but not in IBD or IBS (Priya et al., 2022), leaving open the question of whether such associations are detectable outside of an acute inflammatory context. Our macroscopically non-inflamed samples provide an opportunity to examine this in tissue where active disease is not the dominant signal. We found that transcriptomic and microbiome profiles were significantly associated, demonstrating that there are broad, global similarities between the host transcriptome and mucosal microbiome communities (p = 1×10^-5^, rho = 0.75, Procrustes analysis; **Figure 4B and 4C**).

**Figure 4:**
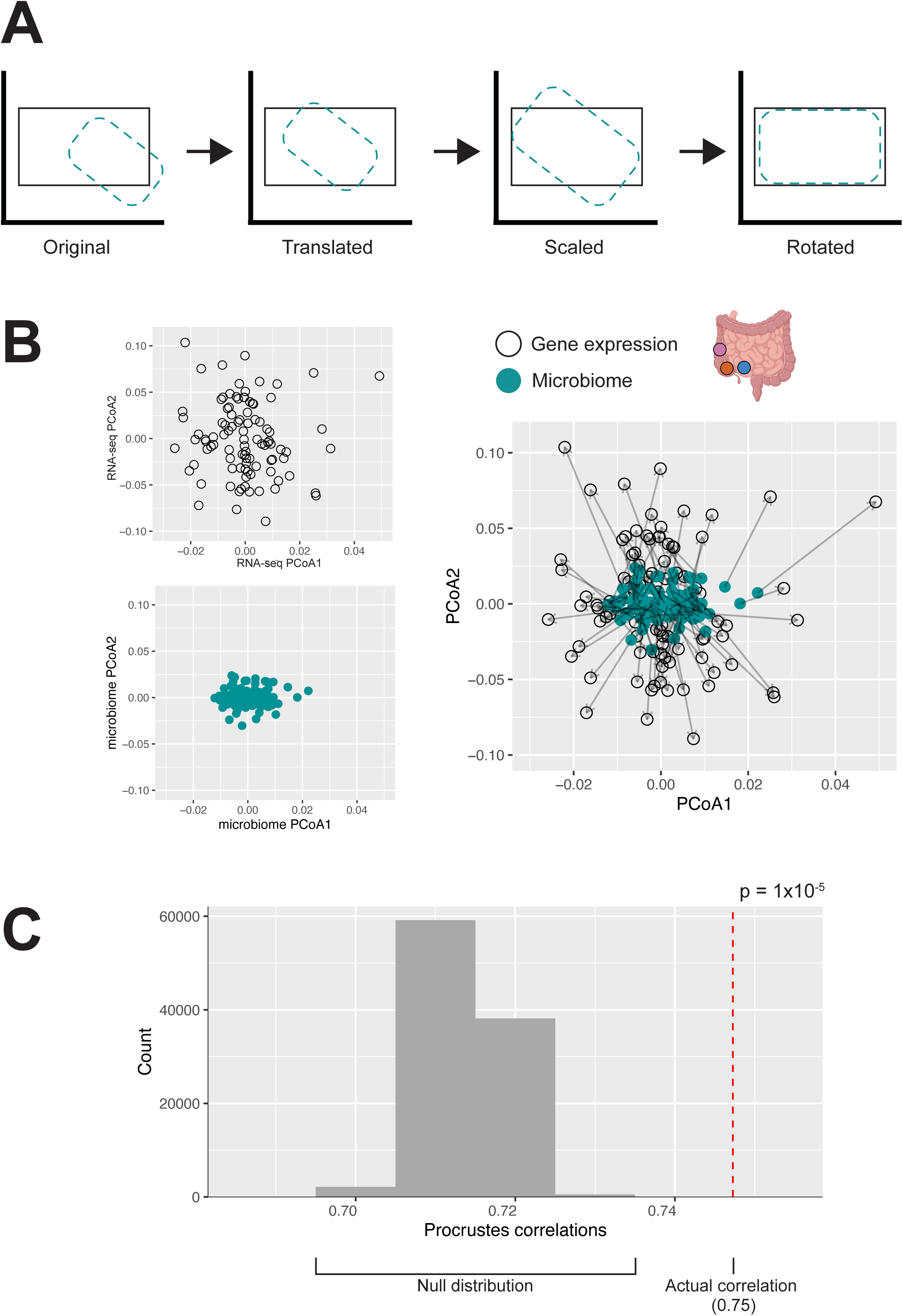
The host transcriptome is significantly globally associated with the microbiome intestine-wide. A) Procrustes analysis superimposes two ordination configurations to assess concordance, here the host transcriptome (black outline) and microbiome (teal dashed outline) distance matrices, by sequentially translating, scaling, and rotating one configuration to maximize alignment with the other. The Procrustes correlations summarize the similarity after superimposition. B) The host transcriptome and microbiome show significant concordance (p = 1×10^-5^, rho = 0.75, Procrustes analysis). The PCoA plots show individuals projected based on Aitchison distances of the RNA-seq data (*top left*, unfilled black circles), Aitchison distances of the microbiome data (*bottom left*, teal), and the two data types superimposed (*right*). Each point represents a sample, with the corresponding microbiome and RNA-seq data connected with an arrow. C) Distribution of Procrustes correlations via permutation test. Sample labels were randomly permuted 99,999 times to generate the null distribution (gray) and compared to the observed Procrustes correlation value (red dotted vertical line) to assess significance.

To gain insights into the physiology underlying this association, we applied sparse canonical correlation analysis (CCA) to identify associations between groups of genes and microbes (**Figure 5A**). Similar to the global association, associations between gene expression-microbe modules have been observed (Hu et al., 2024; Priya et al., 2022). Specifically, genes enriched in pathways for intestinal barrier integrity and the immune system have previously been found to be associated with taxa implicated in IBD. We sought to determine whether similar pathways or types of microbes would be implicated in our study, given that our samples represent a non-acute inflammatory context. Among the top 10 sparse CCA components, components 1, 2, 3, 4, 6, and 10 showed significant correlations between the transcriptomic and microbial canonical variates after multiple test correction (adjusted p < 0.1, rho > 0.49, Pearson correlation; **Supplemental Figure 3, Supplemental Table 3**), revealing multiple modules of co-abundant genes and microbes.

**Figure 5:**
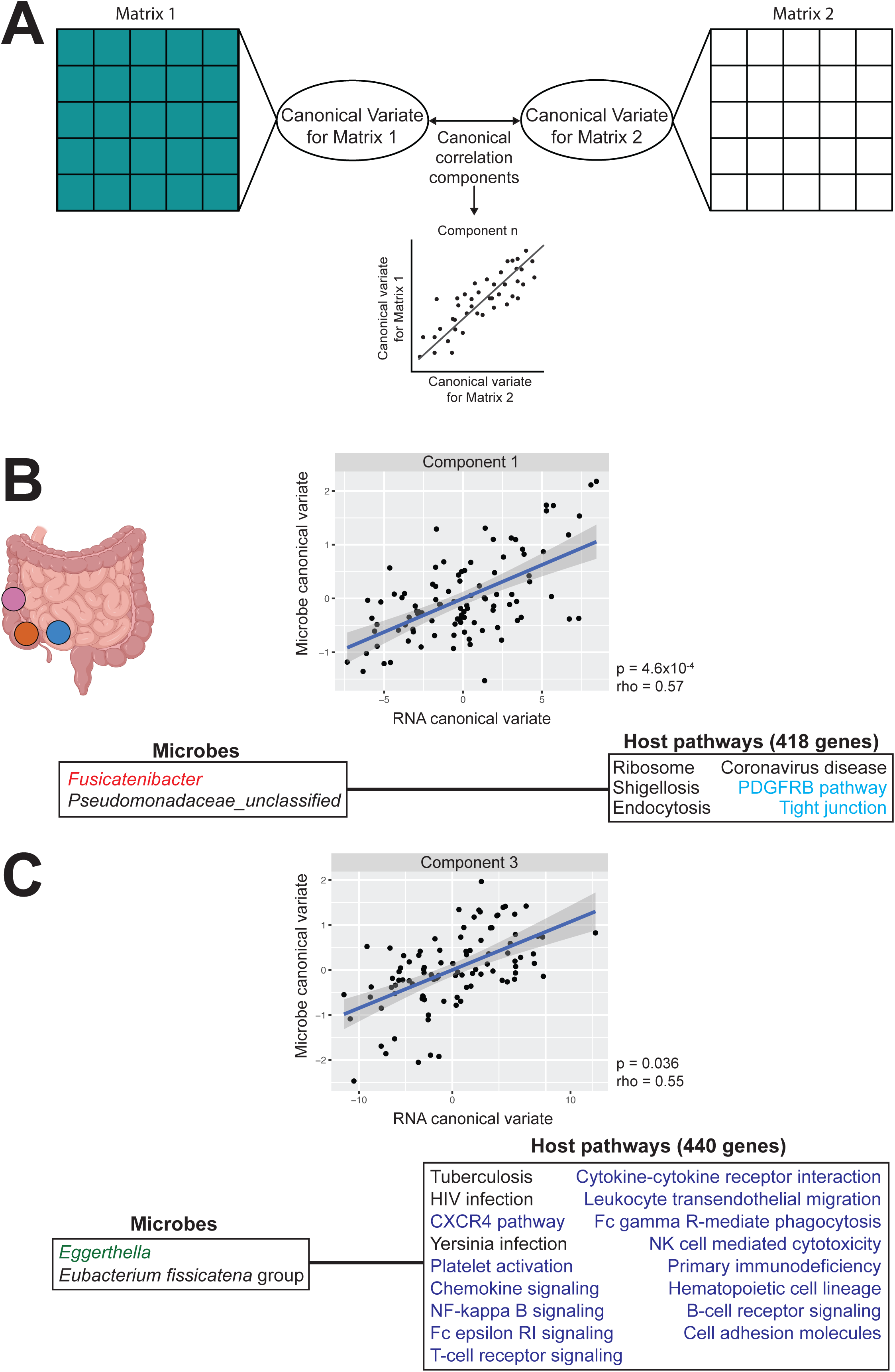
Gut mucosal microbes are associated with the expression of genes enriched for intestinal barrier integrity, immune system, and signaling molecules pathways. A) Sparse canonical correlation analysis (sparse CCA) identifies linear combinations of features from two high-dimensional matrices, here the microbiome (Matrix 1, teal) and host transcriptome (Matrix 2, black), that are maximally correlated with one another. Each canonical correlation component pairs a canonical variate from each matrix, and the correlation between paired variates is assessed for each component. B) Sparse CCA Component 1 for the intestine-wide analysis consists of 2 microbes and 418 host genes enriched in 6 pathways (adjusted p < 0.1, Fisher’s exact test). Heritable microbe *Fusicatenibacter* is colored in red and host pathways for intestinal barrier integrity are colored in cyan. C) Sparse CCA Component 3 for the intestine-wide analysis consisted of 2 microbes and 440 genes enriched in 21 pathways (adjusted p < 0.1, Fisher’s exact test). Opportunistic pathogen *Eggerthella* is colored in green. Host immune and signaling molecule pathways are colored in navy.

To identify the host pathways associated with the microbiome, we performed pathway enrichment analysis on the genes contributing to significant sparse CCA components (**Supplemental Table 4**). Components 1 and 3 were of particular interest, as component 1 includes a microbe from a heritable family and component 3 contains the most enriched pathways at 21 (**Figure 5B and 5C**). Component 1 consists of 2 microbes and 418 genes (**Figure 5B, Supplemental Table 3**). The associated microbes include *Fusicatenibacter,* which is from the heritable Lachnospiraceae family (Goodrich et al., 2016). This component is enriched for six pathways, including the PDGRFB signaling and tight junction pathways (adjusted p < 0.1, Fisher’s exact test; **Supplemental Table 4**). Enrichment in the tight junction pathway is noteworthy, as genes enriched in pathways for gut barrier function have previously been associated with the microbiome (Abdulqadir et al., 2026; Priya et al., 2022; Scott et al., 2020; Ulluwishewa et al., 2011). These associations provide greater insight into microbiome-gut barrier relationships outside of acute inflammatory contexts.

Component 3 consisted of 2 microbes and 440 genes (**Figure 5C, Supplemental Table 3**). The associated microbes included *Eggerthella*, a common gut microbe that is often implicated in gut infections and may promote intestinal inflammation (Gardiner et al., 2015; Shin et al., 2025).

Genes in this component were enriched in the cell adhesion pathway, as well as several immune-related pathways. Specifically, we observed enrichment in the NF-κB pathway, a pathway that plays a crucial role in regulating inflammatory responses (T. Liu et al., 2017; Tak & Firestein, 2001). This pathway activates cell adhesion molecules, which are enriched in this component (adjusted p < 0.1, Fisher’s exact test; **Supplemental Table 4**). This component is also highly enriched in immune pathways, including T-cell and B-cell receptor signaling pathways, which are known to activate the NF-κB pathway. Immune-related genes have consistently emerged as being associated with the gut microbiome, as the intestine needs to maintain adequate defense against microbes and the environment, while also being permeable enough for nutrient absorption (Mowat & Agace, 2014; Zheng et al., 2020). Altogether, our findings suggest that gut microbes associate with the host via immune pathways and barrier pathways even outside of an acute inflammatory context.

We also wished to examine the specific gene-microbe pairs in order to potentially characterize the physiology underlying the relationship. To do this, we implemented elastic net regression, specifically by modeling each microbe against all potential genes. Elastic net is a regularized regression method that simultaneously performs variable selection and handles multicollinearity among predictors, both common features of transcriptomic data, enabling identification of specific gene predictors of microbial abundance from among thousands of candidates. 128 microbes were modeled against 11,316 genes. In total, we identified 13,302 gene-microbe pairs across 6,977 unique genes and 76 unique microbes (**Supplemental Table 5**).

### The terminal ileum is a region of distinct activity between gene expression and microbes

While the analyses above characterized associations between host gene expression and the microbiome that were consistent across gut regions, we also sought to identify relationships that were specific to individual intestinal locations. As with the intestine-wide analysis above, we used a three-pronged approach, similar to Priya et al. (Priya et al., 2022). First, we used Procrustes analysis to examine the global relationships between the transcriptome and microbiome within each region separately. While we observed a significant association between microbiome composition and gene expression across the intestine, when examining by region, we only observed a significant association in the cecum (p = 4.8×10^-4^, rho = 0.69, Procrustes analysis; **Figure 6B**), with no significant associations observed in either the terminal ileum or right colon (p > 0.05 for both, terminal ileum rho = 0.66, right colon rho = 0.63, Procrustes analysis, **Figures 6A and 6C**). The lack of significant associations in the terminal ileum and right colon may reflect limited statistical power, as the rho estimates were comparable but sample sizes for these sites were slightly lower than for the cecum (cecum n = 35 vs. terminal ileum n = 32 and right colon n = 30).

**Figure 6:**
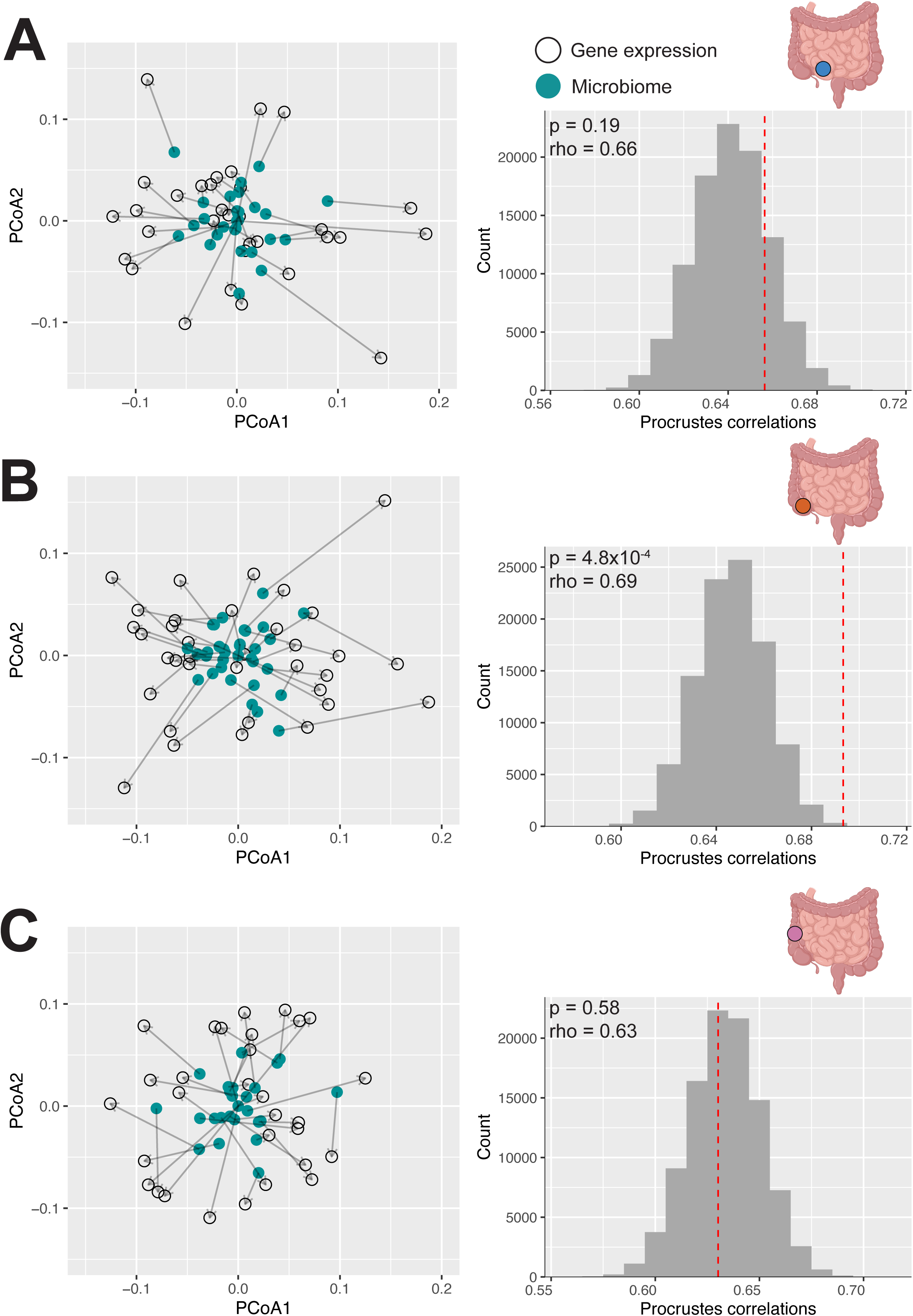
A significant global association between the host transcriptome and microbiome is only observed in the cecum. For each intestinal site, the left panel shows the Procrustes superimposition with arrows connecting the microbiome (teal) and RNA-seq data (black). The right panel shows the permutation distribution, where sample labels were randomly permuted 99,999 times to generate the null distribution (gray) and compared to the observed Procrustes correlation value (red dotted vertical line) to assess significance. A) The host transcriptome and microbiome in the terminal ileum are not globally correlated (p = 0.19, rho = 0.64, Procrustes analysis). B) There was significant concordance between the host transcriptome and the microbiome in the cecum (p = 4.8×10^-4^, rho = 0.69, Procrustes analysis). C) The host transcriptome and microbiome in the right colon are not globally correlated (p = 0.58, rho = 0.63, Procrustes analysis).

Although global associations were not observed in the terminal ileum or right colon, subsets of genes and microbes could show associations. Thus, we performed sparse CCA for each region. We observed significant components for both the terminal ileum and the cecum: components 1, 2, 3, and 9 for the terminal ileum and components 1 and 6 for the cecum (adjusted p < 0.1, rho > 0.71, Pearson correlation; **Supplemental Figure 4, Supplemental Table 6, Supplemental Table 7**). Several of the microbes contributing to the terminal ileum components were from heritable families, specifically Peptostreptococcaceae and Ruminococcaceae, whereas *Desulfovibrio* was the only heritable microbe contributing to the cecum components (Goodrich et al., 2016). No significant components were identified for the right colon, although components 1 and 4 were nominally significant (nominal p < 0.1, rho > 0.76, Pearson correlation; **Supplemental Figure 4, Supplemental Table 8**).

While gene expression-microbe associations have been previously observed in the colon (Hu et al., 2024; Kim et al., 2022; Lloyd-Price et al., 2019; Priya et al., 2022), we were particularly interested in the significantly associated components identified in the terminal ileum, as this site is more difficult to access and therefore understudied. To better understand the physiology underlying these findings, we conducted pathway enrichment of the terminal ileum components (**Figure 7, Supplemental Table 9**). We focused on components 2 and 9, as they consisted of microbes from heritable families. Component 2 of the terminal ileum consists of 2 microbes and 1,724 genes (**Figure 7A, Supplemental Table 6**). This component includes *Peptostreptococcus,* which is part of the heritable family Peptostreptococcaceae (Goodrich et al., 2016) previously associated with genes enriched in intestinal inflammation in gastric disease pathways (Priya et al., 2022). Genes were enriched in intestinal barrier function and repair-related pathways, including the ErbB1 downstream, CDC42 signaling, syndecan-4 signaling, and angiopoietin receptor pathways (adjusted p < 0.1, Fisher’s exact test; **Supplemental Table 9**). Pathways for gut barrier function have been associated with the microbiome across many contexts (Priya et al., 2022) and growth and repair pathways are connected to barrier maintenance (Fink & Wrana, 2023), suggesting a potential microbiome-mediated mechanism for intestinal barrier integrity.

**Figure 7:**
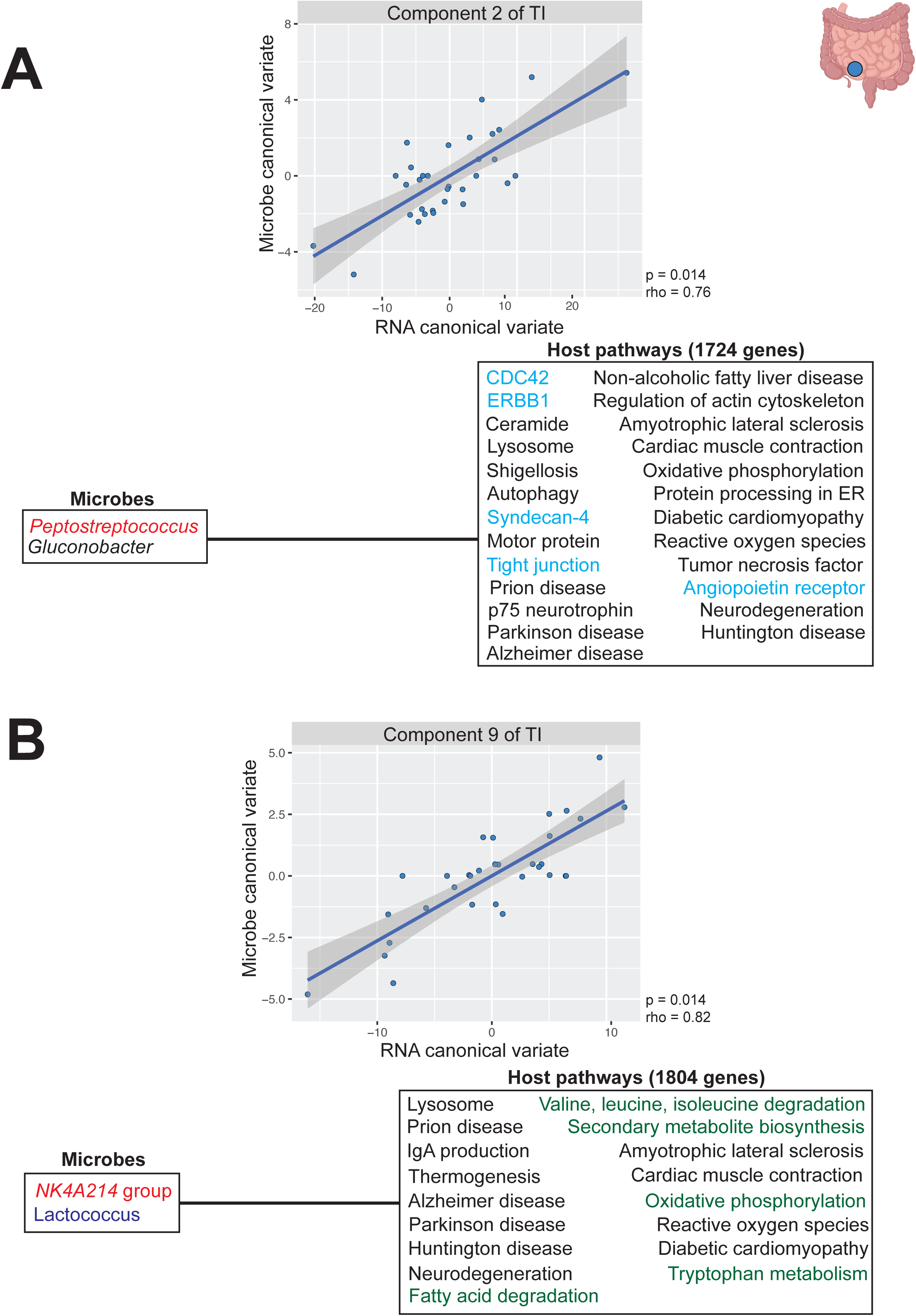
Heritable microbes are associated with genes enriched in intestinal barrier repair and metabolism in the terminal ileum. A) Sparse CCA component 2 in the terminal ileum consisted of 2 microbes and 1724 host genes enriched in 25 pathways (adjusted p < 0.1, Fisher’s exact test). Heritable microbe *Peptostreptococcus* is colored in red and host pathways for intestinal barrier repair are colored in cyan. B) Sparse CCA component 9 in the terminal ileum consisted of 2 microbes and 1804 genes enriched in 17 pathways (adjusted p < 0.1, Fisher’s exact test). Heritable microbe *NK4A214 group* is colored in red, lactic acid bacterium *Lactococcus* is colored in navy, and host pathways for metabolism are colored in green.

Component 9 of the terminal ileum consisted of 2 microbes and 1,804 genes (**Figure 7B, Supplemental Table 6**). This component contained the NK4A214 group microbe, which is part of the heritable family Ruminococcaceae (Goodrich et al., 2016), and the lactic acid bacterium Lactococcus (Dworkin et al., 2006; Onyeaka & Nwabor, 2022). In contrast to the intestine-wide analysis, this component was enriched in several metabolic pathways, which mirrors the roles of the terminal ileum in digestion and nutrient absorption (Mowat & Agace, 2014). Specifically, we observed significant enrichments in amino acid and fatty acid metabolism pathways (adjusted p < 0.1, Fisher’s exact test; **Supplemental Table 9**). We also observed significant enrichment in oxidative phosphorylation, which has previously been associated with the microbiome (Priya et al., 2022). The microbiome contributes broadly to host metabolism by aiding in carbohydrate, vitamin, and protein digestion and also producing key metabolites like short chain fatty acids (SCFAs) (Fan & Pedersen, 2021; Oliphant & Allen-Vercoe, 2019; Rowland et al., 2018). The enrichment of these metabolic pathways with Ruminococcaceae and Lactococcus suggests that microbiome-host metabolic interactions may be particularly prominent in the terminal ileum, consistent with its established role in nutrient absorption.

To further refine our understanding of specific host genes and microbes that are associated within each region, we implemented elastic net regression within each regional dataset. We observed 1,567 gene-microbe pairs across 1,047 genes and 62 microbes within the terminal ileum, 2,305 gene-microbe pairs across 1,549 genes and 69 microbes within the cecum, and 1,455 gene-microbe pairs across 1,242 genes and 43 microbes within the right colon (**Supplemental Table 5**). Notable gene-microbe pairs identified only in the terminal ileum include *PCDH20* and *Faecalitalea* (p = 1.1×10^-4^, rho = 0.63, Pearson correlation; **Figure 8A**) and *ACAT1* and *Lactococcus* (p = 3.9×10^-6^, rho = 0.72, Pearson correlation; **Figure 8B**), the biological implications of which we discuss below.

**Figure 8:**
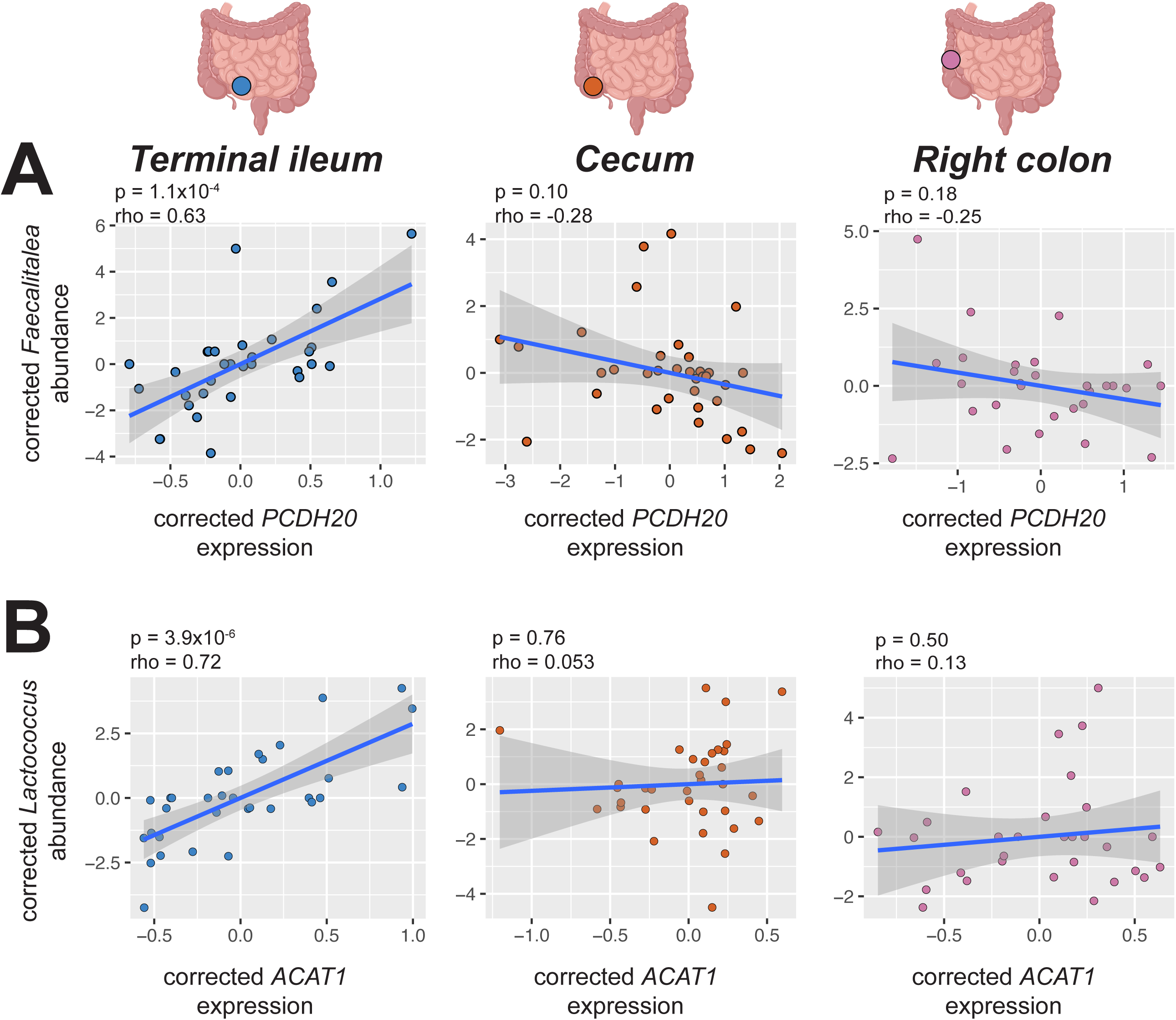
*PCDH20* and *ACAT1* show terminal ileum-specific associations with *Faecalitalea* and *Lactococcus*, respectively. A) Protocadherin 20 (*PCDH20*) and *Faecalitalea* were significantly positively correlated in the terminal ileum (p = 1.1×10^-4^, rho = 0.63, Pearson correlation; left), and not in the cecum (p = 0.10, rho = -0.28; middle) or right colon (p = 0.18, rho = -0.25; right). B) Acetyl-CoA Acetyltransferase 1 (*ACAT1*) and *Lactococcus* were significantly positively correlated in the terminal ileum (p = 3.9×10-6, rho = 0.72, Pearson correlation; left), and not in the cecum (p = 0.76, rho = 0.053; middle) or right colon (p = 0.50, rho = 0.13; right).

## Discussion

Investigations of host gene-microbiome relationships have evolved since the identification of heritable taxa a decade ago. Initial genome-wide association studies of the gut microbiome using fecal samples identified heritable taxa and implicated immune and metabolic genes in shaping microbial composition (Bonder et al., 2016; Davenport et al., 2015; Goodrich et al., 2016; Lim et al., 2017). Subsequent studies leveraging tissue biopsies and gene expression data have begun to characterize the genes and microbes most likely to be physiologically interacting, though these investigations were conducted primarily in disease contexts and within the colon (Hu et al., 2024; Kim et al., 2022; Lloyd-Price et al., 2019; Priya et al., 2022). Here we present a multi-site characterization of mucosal host gene expression-microbiome relationships across three intestinal sites—terminal ileum, cecum, and right colon—using macroscopically non-inflamed tissue to minimize the confounding effects of active disease. We demonstrate that immune and barrier integrity genes are broadly associated with the mucosal microbiome across intestinal sites, patterns that are consistent with prior host genetic studies of the microbiome.

Furthermore, we identify the terminal ileum as a distinct site of gene expression-microbe activity, with region-specific associations involving intestinal barrier maintenance, immune defense, and host metabolic pathways, findings that highlight the importance of including this undersampled region in future investigations.

Across the intestine, genes associated with the mucosal microbiome were enriched in immune-related pathways, consistent with the established role of host immunity in shaping microbial composition and with patterns observed in prior host genetic and transcriptomic studies of the microbiome (Blekhman et al., 2015; Bubier et al., 2021; Priya et al., 2022; Zheng et al., 2020). Among the significant components, sparse CCA component 3 identified a module of genes enriched in the NF-κB signaling pathway associated with *Eggerthella*, a gut microbe implicated in intestinal infections and reported to promote mucosal inflammation through the production of pro-inflammatory metabolites that upregulate TNF-α and IL-6 (Shin et al., 2025). Given that TNF-α and IL-6 are themselves regulators of NF-κB activity (Cabrera-Rivera et al., 2022; Tak & Firestein, 2001), this association may reflect a feedback relationship between *Eggerthella*-derived inflammatory signals and host NF-κB-mediated immune responses. The fact that this pattern is detectable in macroscopically non-inflamed tissue suggests that low-level immune-microbe interactions along this axis may be a feature of baseline intestinal physiology rather than being exclusive to active disease states, though this interpretation requires direct experimental validation.

Genes associated with the mucosal microbiome across intestinal sites were enriched for intestinal barrier maintenance pathways, specifically tight junctions. Furthermore, genes enriched in barrier integrity pathways have previously been associated with the microbiome in IBD contexts (Priya et al., 2022), and our findings extend these observations to macroscopically non-inflamed tissue. Interestingly, the tight junction gene module was associated with *Fusicatenibacter*, a heritable SCFA-producing member of Lachnospiraceae found in lower abundance in the feces of IBD patients (Aoyama et al., 2025; Blesl et al., 2025; Takeshita et al., 2016). *Fusicatenibacter* produces multiple SCFAs including acetate, lactate, formate, propionate, and succinate (Jin et al., 2019; Medawar et al., 2021; Pihelgas et al., 2024; Takada et al., 2013), metabolites that are well-established modulators of tight junction expression and epithelial barrier function (Yue et al., 2022). Together, these findings are consistent with a model in which variation in SCFA-producing taxa such as *Fusicatenibacter* co-occurs with host transcriptional programs involved in epithelial barrier maintenance, suggesting a potential interface through which host genetics and microbial metabolites may jointly influence mucosal integrity.

While intestine-wide analyses identified barrier-related gene modules associated with the mucosal microbiome, region-specific analyses revealed additional nuance in the terminal ileum. We observed a positive association between protocadherin 20 (*PCDH20*) and *Faecalitalea* exclusively in the terminal ileum, with no comparable relationship detected in the cecum or right colon. *PCDH20* encodes a cadherin protein with established roles in intestinal epithelial morphology and microbial balance (Huang et al., 2023). Its deletion in mice disrupts gut microbiota composition and impairs intestinal barrier function under inflammatory conditions.

*PCDH20* is additionally necessary for tuft cell microvilli formation (Ankenbauer et al., 2026). Stimulation of tuft cells by microbially produced metabolites can result in type 2 immune responses, triggering antimicrobial peptide release that can shape microbial composition (Fung et al., 2023). *Faecalitalea* is an SCFA-producing gut microbe (De Maesschalck et al., 2014; Ma et al., 2020), and SCFAs are known for their positive role in intestinal barrier function (P. Liu et al., 2021; Parada Venegas et al., 2019; Pérez-Reytor et al., 2021). Together, these observations raise the possibility that SCFA production by *Faecalitalea* may influence barrier-related gene expression in the terminal ileum, though the precise mechanism and directionality of this relationship remain to be established.

Beyond individual gene-microbe associations, the functional categories of host genes associated with the mucosal microbiome also differ by intestinal site. Genes enriched in metabolic pathways, including amino acid metabolism, were associated with the microbiome uniquely in the terminal ileum and not in the cecum or right colon, consistent with the established role of the terminal ileum in nutrient absorption (Mowat & Agace, 2014). A notable example is the positive association between acetyl-CoA acetyltransferase (*ACAT1*) and *Lactococcus* in the terminal ileum. *ACAT1* is involved in various metabolic processes including fatty acid metabolism and ketogenesis (Goudarzi, 2019), and *Lactococcus* is a lactic acid bacterium whose fermentation products include lactate and SCFAs (Dworkin et al., 2006; H. Tang et al., 2023). *Lactococcus* has also been shown to affect host metabolic processes, including lipid metabolism and glucose metabolism (M. Wang et al., 2024; Yoda et al., 2026), raising the possibility that microbially-produced metabolites, including lactate, may intersect with the ACAT1-mediated metabolic pathways in the intestinal epithelium. The precise mechanism underlying this association remains to be established, but the enrichment of metabolic pathway associations in the terminal ileum suggests this undersampled site may be a particularly important locus of host-microbiome metabolic interactions.

The functional landscape of genes associated with the mucosal microbiome in our study was consistent with patterns emerging from microbiome genome-wide association studies. Host genetic variants most robustly associated with gut microbiome composition have clustered in or near genes related to immune defense, mucosal barrier integrity, and metabolic sensing (Bonder et al., 2016; Goodrich et al., 2016; Lopera-Maya et al., 2022). Our transcriptomic findings recapitulated this same functional landscape: immune signaling genes including those enriched in NF-κB pathways, barrier integrity genes including tight junction components, and metabolic genes including those involved in fatty acid oxidation and ketone body metabolism are among the prominent associations we identify (**Supplemental Table 4** and **Supplemental Table 6**). Direct gene-level overlaps between our elastic net results and published microbiome genome-wide association study (GWAS) loci include *PTPRG* (Kurilshikov et al., 2021), a protein tyrosine phosphatase that regulates cell growth and differentiation (Boni et al., 2022), and *ADCYAP1* (Kurilshikov et al., 2021), which encodes a neuropeptide (PACAP) with roles in gut motility and gastric acid secretion (Fujimiya & Inui, 2000; Oh et al., 2005), as well as thematic overlap in heparan sulfate sulfotransferase biology (*HS3ST1* in our cecum results; *HS3ST4* at a GWAS locus for Faecalibacterium) (Ishida et al., 2020). This convergence across genetic and transcriptomic levels of analysis is consistent with the framing of gene expression as a dynamic intermediate between host genomic variation and microbial community composition and suggests that the biological axes through which host genetics shapes the microbiome are similarly detectable at the transcriptional level in mucosal tissue.

While we demonstrate gene expression-microbe associations both intestine-wide and within regions, there are several limitations to our study. First, our power to detect associations is limited by our modest sample size. Acquiring sufficient sample size from surgically resected tissue is inherently difficult, and we chose to limit our investigation to a controlled cohort to minimize confounders rather than pooling across disease contexts, inflammation statuses, or separate cohorts as some prior studies have done. As a result, we were likely underpowered to detect some associations with small effect sizes. The absence of significant modular associations in the cecum, for example, may reflect limited power rather than lack of true signal, given that we did observe a significant global association there and because the cecum is known for its microbial and host immune activity (James et al., 2020; Yang et al., 2025). In another example, site-specific Procrustes analysis only reached significance in the cecum where the sample size was largest despite similar rho values in the terminal ileum and right colon comparisons, which may reflect limited power rather than a true absence of signal. Thus, our results should be considered in the context of our limited sample size.

Second, while surgical resection provided access to otherwise unobtainable mucosal tissue, the patients from whom the samples were collected underwent surgical preparation. This process generally entails taking antibiotics and bowel preparation, both of which may alter microbial composition and gene expression relative to a non-prepped state (Ferrie et al., 2021; O’Brien et al., 2013). As a result, these methodological constraints inherent to working with tissue from these anatomical sites in humans should be considered when interpreting our findings.

Finally, we cannot determine whether identified relationships are causal or bidirectional. Either the microbe could be regulating gene expression, or host gene expression could be shaping microbial relative abundance. Additionally, we cannot exclude the possibility that both are driven by unmeasured confounders. Future work should prioritize identifying the directionality of these associations and characterizing underlying mechanistic pathways. This will be crucial for gaining an understanding of host-microbiome crosstalk in health and disease and potentially targeting elements for therapeutic benefit.

## Conclusions

Here, we demonstrated that there are significant associations between host gene expression and the gut microbiome both across intestinal sites and within specific regions. Immune and barrier integrity gene modules were broadly associated with the mucosal microbiome, consistent with patterns emerging from host genetic studies of the microbiome. The terminal ileum emerged as a particularly notable site of host-microbiome interaction, with region-specific associations involving metabolic and barrier-related pathways that highlight the value of including this undersampled region in future investigations. Together, these findings advance our understanding of the physiological axes through which host gene expression and the mucosal microbiome are linked, and provide a foundation for future work aimed at uncovering the mechanistic basis of host-microbiome crosstalk in intestinal health and disease.

## Methods

### Ethics statement

Sample collection for the Carlino Family Inflammatory Bowel and Colorectal Diseases Biobank was approved by the Penn State Milton S. Hershey Medical Center Institutional Review Board (IRB Protocol: PRAMSHY98-057), with subsequent genomic data collection approved by the Pennsylvania State University Institutional Review Board (IRB Protocol: STUDY00020731). All specimens were collected with informed consent between 2015-2021.

### Sample collection

All samples were collected and stored by the Carlino Family Inflammatory Bowel and Colorectal Disease Biobank at the Penn State Milton S. Hershey Medical Center (Mankarious et al., 2023), having been collected during colectomy under informed consent. We selected samples from three locations—terminal ileum, cecum, and right colon—to evaluate host transcriptome-microbiome relationships across the gut. To minimize the confounding effects of active inflammation, only samples from macroscopically non-inflamed sites determined via visual inspection at the time of surgery were selected, and only individuals with paired full-thickness tissue resections and mucosal scrapings were retained. Mucosal scrapings were collected by scraping a microscope slide against the intestinal lining. Most full-thickness tissues were stored in RNAlater, although one sample was flash frozen. Both sample types were immediately preserved in cryogenic storage tubes at -80°C for long-term storage in the biobank. Samples were shipped overnight to The Pennsylvania State University - University Park (PSU) on dry ice and immediately stored at -80°C upon arrival.

Mucosal scrapings were selected for microbiome characterization to reduce host contamination during extraction (**Supplemental Table 10**) and corresponding full-thickness tissues were selected for RNA extraction (**Supplemental Table 11**). In total, 159 mucosal scrapings were collected for microbiome analysis from 102 Crohn’s disease patients: 58 terminal ileum, 55 cecum, and 46 right colon samples. For RNA-seq analysis, a total of 158 full-thickness tissues were collected from 102 Crohn’s disease patients: 58 terminal ileum, 54 cecum, and 46 right colon. An additional 20 right colon mucosal scrapings and 15 full-thickness tissue samples from colorectal cancer patients were included to evaluate the extent to which underlying inflammation from IBD affects microbial and transcriptomic profiles at macroscopically non-inflamed sites.

### DNA extraction and 16S rRNA amplicon sequencing

All pre-PCR steps were conducted in a specialized low biomass, low contamination lab space to reduce potential environmental contamination (Fierer et al., 2025). Total DNA was extracted from mucosal scrapings using the QIAamp DNA Microbiome kit (Qiagen, Germantown, MD).

This kit was selected for its upstream host tissue-degradation steps, as host carryover was a major concern. We largely followed the manufacturer’s protocol, with the exception of the initial tissue lysis steps. These steps involved adding the mucosal scrapings to 750 µl of DNA Elution buffer in a ZR BashingBead 2.0 mm Lysis Tube (Zymo Research, Irvine, CA), and then bead beating the tissue at maximum speed for 1 minute using the Bead Ruptor 96 Well Plate Homogenizer (Omni International, Kennesaw, GA). Negative controls were included at the beginning, middle, and end of each extraction batch to track potential kit contamination, environmental contamination, and cross contamination. Quantitation on the Qubit Fluorometer confirmed negligible DNA concentrations (**Supplemental Table 10**), indicating the absence of large-scale contamination events during extraction.

The V1-V2 hypervariable region of the 16S rRNA gene was PCR amplified for all DNA extracts and negative controls. This region of the 16S rRNA gene was selected for its reduced off-target amplification of host DNA (Walker et al., 2020). Standard PCR protocols and cycle parameters were followed (Caporaso et al., 2012), with all samples and controls amplified in triplicate using 35 cycles of PCR to ensure that sufficient material was recovered from these low biomass samples. PCR negative controls were included at the beginning and end of each PCR batch to track potential contamination sources, as well as DNA positive controls (ZymoBIOMICS Microbial Community DNA Standard, Zymo Research, Irvine, CA).

To avoid sequencing off-target amplicons, the QIAquick Gel Extraction Kit was used to specifically target the expected microbial amplicon length of 310 bp (Qiagen, Germantown, Maryland). The gel extracts were cleaned using the AxyPrep MAG PCR Clean-up kit to remove any remaining salts (Corning, Corning, NY). Final DNA concentrations were measured using the Qubit dsDNA Quantification Assay Kit (Invitrogen, Waltham, MA). Cleaned amplicons were subsequently submitted to the PSU Huck Institutes of the Life Sciences Genomics Core Facility for library preparation. Sample concentrations were normalized using the SequalPrep Normalization Plate Kit (Thermo Fisher Scientific, Waltham, MA) and subsequently sequenced 300 x 300 paired-end on the Illumina NextSeq 2000 using the P1 600 cycle kit (Illumina, San Diego, CA). Notably, the SequalPrep library normalization procedure preferentially enriches low-concentration samples, such that negative controls yielded detectable read counts upon sequencing, despite having negligible DNA concentrations at quantification. This is an expected consequence of the normalization procedure and does not reflect a contamination event.

Rather, these negative control libraries were used to identify and remove contaminant amplicon sequence variants (ASVs).

### Amplicon sequence data quality control and decontamination

Raw sequences were processed using a custom Snakemake (v.8.14.0) pipeline that verified the quality, length, and depth of the reads using FastQC v.0.12.1 and filtered any remaining host contaminant reads (Andrews, 2010; Bush et al., 2020; Mölder et al., 2021). Specifically, we followed the two-step host read removal pipeline outlined by Bush et al. (Bush et al., 2020). In brief, reads were aligned to the human reference genome (GRCh38) using bowtie2 v.2.5.2 (Langmead & Salzberg, 2012). Reads that did not map to the human genome via bowtie2 were extracted and subsequently aligned to the human genome again, this time with SNAP v.2.0.3 (Zaharia et al., 2011). Reads that did not map to the human genome via both bowtie2 and SNAP were considered microbial and retained for further processing (**Supplemental Figure 5)**. Samples with very low read depth (< 10 reads) were removed (n = 2).

Microbial reads underwent cleaning and processing using QIIME 2 2024.2 (Bolyen et al., 2019). First, primers were trimmed using cutadapt via the q2–cutadapt plugin (Martin, 2011).

Untrimmed reads or reads < 100 bp were removed. Trimmed paired-end reads were merged using VSEARCH via q2–vsearch (Rognes et al., 2016). Merged reads with ambiguous base calls or bases below a quality score of 4 were removed via q2–quality-filter (Bokulich et al., 2013). Merged filtered reads were denoised using deblur via q2–deblur (Amir et al., 2017) and trimmed to 270 bases based on expected read length. Amplicon sequence variants (ASVs) were aligned with mafft and then used to infer a phylogenetic tree with fasttree2 via q2–phylogeny (Katoh et al., 2002; Price et al., 2010). ASVs were classified using a naïve Bayes taxonomy classifier via q2-feature-classifier (Bokulich et al., 2018). This classifier was trained by matching Silva 138 99% reference sequences to the amplicon region based on our primer set, trimming the expected region length of 270 bases, and then training the classifier using these optimized reference sequences (Quast et al., 2013). The resulting feature tables, taxonomic assignments, phylogenetic tree, and metadata were exported from QIIME 2 into R version 4.4.3 using qiime2R v.0.99.6 (Bisanz, 2018). Singletons were removed and ASVs were subsequently analyzed in phyloseq v.1.50.0 for further decontamination and downstream analysis (McMurdie & Holmes, 2013).

Next, samples were examined for additional potential contamination. Due to the low-biomass nature of the samples, contamination is a major consideration, especially due to varying starting sample mass (0.1mg – 80mg) (Fierer et al., 2025). Although negative control sample DNA concentrations were negligible after DNA extraction and library preparation which indicates that a large-scale contamination event was unlikely to have occurred, contamination from the environment, from kits, or cross-sample contamination could have still occurred to a lesser extent. To document and control for potential contamination, several bioinformatic measures were taken. First, any samples with library concentrations lower than those of the negative controls in the same batch were removed (n = 14 removed). In addition, samples with tissue mass < 0.5mg were removed from the analysis (n = 8). To account for environmental contamination, potential contaminants were identified and removed via decontam v.1.26.0 using the frequency method, thereby removing 168 ASVs (Davis et al., 2018). Next, singletons and any taxa that did not appear with at least 5 reads across at least two samples were removed to account for potential contaminants or sequencing artifacts. Following these procedures, negative control samples were significantly different from true samples (p = 1×10^-5^, PERMANOVA; **Supplemental Figure 6**) and positive control samples primarily consisted of the 8 expected taxa in relatively even proportion (**Supplemental Figure 7**), thus demonstrating the ability of our approach to prevent and remove contamination without removing true microbes.

Finally, samples below 65,467 reads were removed in preparation for downstream rarefaction (n = 5). This read depth was chosen based on the minimum read count of samples that were sequenced at sufficient depth while also minimizing sample loss due to low read depth (**Supplemental Figure 8**). Raw counts were used for tools with built-in compositionally-aware transformation methods, such as MaAsLin2 (Mallick et al., 2021). For all other microbiome analyses, counts were transformed to relative abundances using total-sum scaling. In total, 50 terminal ileum, 43 cecum, and 39 right colon samples were used for downstream analysis, as well as the 18 right colon samples from colorectal cancer patients.

### Microbiome covariate correction

Biological and technical confounders have been major considerations for microbiome research, as both shape the microbiome (Y. Wang & LêCao, 2020). As a result, the microbiome data were tested for association with multiple covariates. To do this, Bray-Curtis, unweighted UniFrac, and weighted UniFrac distances were calculated from the microbiome counts using phyloseq v.1.50.0 and rbiom v.2.2.0 packages for Bray-Curtis and UniFrac, respectively (McMurdie & Holmes, 2013; Smith, 2026), and then the resulting distances were transformed to principal coordinates (Bray & Curtis, 1957; Lozupone & Knight, 2005). Principal coordinates were individually tested for association with sex, age, age at time of surgery, library concentration, and extraction batch. From this, sex, age, library concentration, and extraction batch were identified as covariates.

### Diversity analyses

Five metrics were used to examine alpha diversity: Faith’s phylogenetic diversity, Fisher’s alpha, Shannon index, Simpson index, and species richness (Faith, 1992; Fisher et al., 1943; Shannon, 1948; Simpson, 1949). Each metric was calculated by rarefying counts using the rarefy_even_depth function from phyloseq v.1.50.0 (subsampling to 65,467 sequences) and then calculating the alpha diversity metric (McMurdie & Holmes, 2013). This was repeated 1,000 times to account for randomness in rarefaction, and the mean alpha diversity value was calculated. Diversity metric values were corrected for covariates by fitting a linear mixed model using the lmer function from the R package lme4 v.1.1-37 (D. Bates et al., 2025). Specifically, the model included the covariates as predictors, metric values as the response variable, and patient ID as a random effect, and the residuals were used as the corrected diversity metric values. ANOVA tests were conducted to evaluate whether alpha diversity significantly differed based on location.

Beta diversity was evaluated via three methods: Bray-Curtis, unweighted UniFrac, and weighted UniFrac using phyloseq v.1.50.0 and rbiom v.2.2.0 packages for Bray-Curtis and UniFrac, respectively (Bray & Curtis, 1957; Lozupone & Knight, 2005; McMurdie & Holmes, 2013; Smith, 2026). Distances were calculated from relative abundances and then subjected to principal coordinates analysis (PCoA). Covariate correction was performed as described for alpha diversity above, with principal coordinate values included as the response variable in place of diversity metric values.

### Differential abundance analysis

To identify differentially abundant taxa between locations, microbiome counts were agglomerated to the genus level, centered log-ratio (CLR) transformed, and analyzed using MaAslin2 v.1.20.0, which implements a generalized linear mixed model (Mallick et al., 2021). Covariates identified earlier were included in the model, with patient ID included as a random effect. Multiple test correction was conducted via the Benjamini-Hochberg method (Benjamini & Hochberg, 1995). Adjusted p-values < 0.05 were considered significant.

### RNA extraction and RNA-sequencing

Total RNA was extracted from full-thickness tissue resections, most of which were stored in RNAlater, with the exception of one flash-frozen sample. Tissue samples were homogenized using a pestle in 1 ml of TRIzol (Invitrogen, Waltham, MA). Chloroform was added and the samples were centrifuged at 12000 x g for 15 minutes to separate the aqueous and organic phases. The aqueous layer was then collected and purified using the RNeasy Midi Kit (Qiagen, Germantown, MD) and treated with DNase I to remove genomic DNA. RNA concentration was determined via NanoDrop (Thermo Scientific, Waltham, MA), and RNA integrity was evaluated via Agilent Bioanalyzer (Agilent Technologies, Santa Clara, CA). Extracts with RNA integrity number (RIN) < 6 were not processed further and those with RNA concentration < 25 ng/µl were concentrated using a SpeedVac. cDNA libraries were generated using the Illumina Stranded mRNA Prep kit (Illumina, San Diego, CA) and their concentrations were measured via the Bioanalyzer. Libraries with cDNA concentration < 1 ng/µl were excluded from further analysis.

Individual libraries were subsequently submitted to the PSU Huck Institutes of the Life Sciences Genomics Core Facility for library pooling and 100 bp single-end Illumina NextSeq 2000 sequencing using the P4 100 cycle kit (Illumina, San Diego, CA).

### RNA-seq data quality control and cleaning

Raw sequences were processed using a custom Snakemake (v.8.14.0) pipeline that verified the quality, length, and depth of the reads using FastQC v.0.12.1, aligned the reads to the human genome, and generated a genomic count table (Andrews, 2010; Mölder et al., 2021). Reads were aligned to the human reference genome (GRCh38) using STAR v.2.7.11b (Dobin et al., 2013). A genomic feature count table was generated using featureCounts v.2.0.6 (Liao et al., 2014). Lowly expressed genes were filtered by only keeping genes that had at least 10 counts across at least 15 samples (the smallest group in the dataset). Variance-stabilizing transformation was applied to the counts using the vst function in DESeq2 v.1.46.0 (Love et al., 2014). Finally, samples were removed from the analysis due to either poor quality RNA (RIN < 7) or identification of potential sample swaps (n = 14). In total, 39 terminal ileum, 44 cecum, and 37 right colon samples were retained for downstream analysis, along with 15 right colon samples from colorectal cancer patients.

### RNA-seq covariate correction

Similar to the microbiome, gene expression patterns can be confounded by biological and technical factors, and genomic feature counts were tested for potential covariates (Englander, 2005; Leek et al., 2010; Yamamoto et al., 2022). Principal components analysis was applied to gene counts, and principal components (PCs) were individually tested for association with sex, age, age at time of surgery, library concentration, RIN score, extractor, extraction batch, library batch, and sequencing batch. From this, sex, age, library concentration, RIN score, extractor, and sequencing batch were identified as covariates.

### RNA-seq analysis

Differential gene expression analysis was conducted using the dream function in the variancePartition package v.1.36.3 (Hoffman & Roussos, 2020; Hoffman & Schadt, 2016).

RNA-seq counts were first filtered for genes with CPM above 0.5 across at least 37 samples (smallest group size) and then remaining filtered counts were trimmed mean of M-values (TMM) normalized using edgeR v.4.4.2 (Robinson et al., 2010). Covariates identified earlier were included in the model (sex, age, library concentration, RIN score, extractor, and sequencing batch), with patient ID included as a random effect. Multiple test correction was conducted via the Benjamini-Hochberg method (Benjamini & Hochberg, 1995). Genes with log fold changes > 2 and adjusted p-values < 0.05 were considered significant.

### Gene expression-microbiome integration

Associations between gene expression and the microbiome were examined from samples with both data types available (n = 32 terminal ileum, n = 35 cecum, and n = 30 right colon). RNA-seq data were prevalence filtered, with genes retained if they had at least 10 counts across at least 33% of the overlapping samples. The retained genes were normalized using variance stabilizing transformation (VST). Non-protein-coding genes were filtered out.

For cross-location analyses, covariates pertaining to RNA sample processing and patient data were regressed out, along with specimen location. Because some samples across locations were from the same patient, patient ID was also included in the model as a random effect to control for intra-individual variation. Finally, genes in the lowest 25% quantile of variance were filtered out, resulting in a final dataset of 11,316 genes.

For within-region analyses, samples were similarly filtered and normalized, while also accounting for location in the design formula. Samples were then split by location and then corrected for covariates pertaining to data type and patient data for each location separately. Genes in the lowest 25% quantile of variance were filtered out, resulting in a dataset of 10,895 genes per location.

For the microbiome, ASVs were used for global-level analyses and counts were agglomerated to the genus level for group- and individual-level analyses (see below). Counts were transformed to relative abundances. For group- and individual-level analyses, taxa with lower than 0.01% relative abundance across at least 10% of the samples were filtered out and then CLR transformed via the microbiome R package v.1.28.0 (Lahti & Shetty, 2017). No filters were applied to global analyses. Finally, covariates pertaining to microbiome sample processing and patient data were regressed out. Ultimately, this resulted in 128 genera retained for analyses across the gut. When analyzed by region, samples were split prior to transformation and processed separately. The resulting datasets included 129 genera in the terminal ileum, 122 genera in the cecum, and 141 genera in the right colon.

Gene expression-microbiome relationships were evaluated using a three-level approach, termed “global”, “group-group”, and “individual”. Specifically, “global” analyses assessed broad compositional similarities between the transcriptome and microbiome across host genes and taxa. “Group-group” analyses identified groups of genes associated with groups of microbes, motivated biologically as genes within pathways often are co-regulated and microbes frequently form co-abundance groups. “Individual” analyses identified relationships between specific host genes and microbes. Because relationships may exist at any of these levels, a multi-level approach was necessary for comprehensive characterization of host-microbiome interactions.

To identify “global” relationships between host gene expression and microbiome composition, Procrustes analysis was used via vegan v.2.6-10 (Dixon, 2003; Gower, 1975). Aitchison distance was calculated between samples after adding a pseudocount of 0.000001 and then visualized using PCoA (Aitchison, 1982; Aitchison et al., 2000). The resulting matrices were corrected for corresponding covariates and Procrustes analysis was conducted using the corrected matrices (**Figure 4A**). To check for significance, rows of the microbiome matrix were randomly permuted 99,999 times to generate the null distribution and then compared to the nonpermuted Procrustes correlation value. A global relationship was considered significant at a permutation p < 0.05 threshold.

To identify “group-group” relationships (i.e. whether groups of co-expressed genes are significantly associated with groups of co-abundant genera), sparse CCA was conducted. Sparse CCA is similar to traditional CCA, in that it identifies the canonical components that maximize the correlation between two datasets (**Figure 5A**) (Witten et al., 2009; Witten & Tibshirani, 2009). Traditional CCA, however, cannot be applied to situations in which the number of predictors greatly outweighs the number of samples. Sparse CCA addresses this limitation in high dimensional datasets by identifying the subset of variables (hence the “sparse”) that maximally explains the correlation between two datasets. To do so, it implements a lasso penalty for each dataset to identify the most relevant predictors in an association. To identify the optimal lasso penalty, a grid-search approach with leave-one-out-cross-validation (LOOCV) was implemented. The lasso penalty that resulted in the greatest correlation value was selected for the sparse CCA model using PMA v.1.2-4 as described by Priya et al. (Priya et al., 2022; Witten et al., 2009; Witten & Tibshirani, 2009). The first 10 sparse CCA components were extracted and subsequently tested for significance of the correlation via LOOCV at a p-value cutoff of 0.1. Genes contributing to the components were analyzed for pathway enrichment using the enricher function in clusterprofiler v.4.14.6 (Wu et al., 2021; Yu et al., 2012), with Kyoto Encyclopedia of Genes and Genomes (KEGG) database FTP release 2022–11-07 (Kanehisa & Goto, 2000) and Pathway Interaction Database (PID) (Schaefer et al., 2009) as reference databases. This function conducts over representation analysis and applies a one-sided Fisher’s exact test to identify enriched pathways (Boyle et al., 2004). Multiple test correction was conducted via the Benjamini-Hochberg method (Benjamini & Hochberg, 1995). Pathways were considered significantly enriched if they had an adjusted-p value less than 0.1 and a gene set size greater than or equal to 10.

To identify “individual” relationships between gene expression and microbial relative abundance, elastic net regression was used. Elastic net regression is a type of regression that is optimized for high-dimensional data with multicollinear predictors (Friedman et al., 2010). Specifically, it combines both the power of lasso to filter out irrelevant predictors and also the power of ridge regression to account for multicollinearity between predictors. This method was performed within each region. The model took the following form:

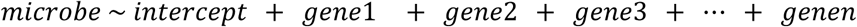

As covariates were already regressed out upstream, they were not included in the model. A model was built and fit for each microbe using packages glmnet v.4.1-8 and caret v.7.0-1 (Friedman et al., 2010; Kuhn, 2008). Specifically, the model was simultaneously tuned for the best alpha and lambda via 5-fold cross validation repeated 5 times. The best lambda was then used to predict the coefficients for the gene predictors. Nonzero coefficients were extracted and used to identify gene expression-microbe associations. Due to the computationally demanding and time-intensive nature of the model tuning process, these steps were conducted in parallel using the packages foreach v.1.5.2 and doParallel v.1.0.17 (Folashade, Ooi, et al., 2022; Folashade, Weston, et al., 2022).

## Data, Metadata, and Code Availability

Microbiome sequence data can be retrieved from the Sequence Read Archive (SRA) under BioProject Number PRJNA1311722. RNA-seq data can be retrieved from the SRA under BioProject Number PRJNA1444093. Full microbiome metadata is available in **Supplemental Table 10** and full gene expression metadata is available in **Supplemental Table 11**. All source code to recreate the analyses in the paper is available at https://github.com/davenport-lab/intestinal_GE_microbiome.

## Acknowledgements

The authors would first and foremost like to thank the anonymous patients who were willing to donate tissue to The Carlino Family Inflammatory Bowel and Colorectal Diseases Biobank that enabled this research to be conducted. Additionally, we would like to acknowledge the Huck Institutes’ Genomics Core Facility (RRID:SCR_023645) for sequencing microbiome and

RNA-seq data, the Genomics Research Incubator (RRID:SCR_024530) for RNA-seq library prep assistance, and the Institute of Computational and Data Sciences (RRID:SCR_026424) for high performance computing resources.

This work was supported by the National Institutes of Health (R35GM146980) to ERD, with EPR supported by the Computation, Bioinformatics, and Statistics training grant (T32GM102057).

The funders had no role in study design, data collection and analysis, decision to publish, or preparation of the manuscript.

## Abbreviations

ANOVA: analysis of variance
ASV: amplicon sequence variant
bp: base pair
CCA: canonical correlation analysis
CLR: centered log-ratio
CPM: counts per million
Faith’s PD: Faith’s phylogenetic diversity
GWAS: genome-wide association study
IBD: inflammatory bowel disease
IBS: irritable bowel syndrome
IRB: Institutional Review Board
logFC: log fold change
LOOCV: leave-one-out cross-validation
PCA: principal component analysis
PCoA: principal coordinate analysis
PCs: principal components
PERMANOVA: permutational multivariate analysis of variance
RIN: RNA integrity number
SCFA: short-chain fatty acid
sparse CCA: sparse canonical correlation analysis
SRA: Sequence Read Archive
TMM: trimmed mean of M-values
VST: variance-stabilizing transformation

## Notes

https://github.com/davenport-lab/intestinal_GE_microbiome

### Competing Interest Statement

The authors have declared no competing interest.

## References

Abdulqadir, R., Al-Sadi, R., Gupta, Y., Rawat, M., & Ma, T. (2026). Probiotic bacteria Bifidobacterium bifidum upregulation of intestinal epithelial tight junction barrier is mediated by TLR-2/TLR-6 receptor complex activation of occludin gene. Npj Biofilms and Microbiomes, 12(1), 37. 10.1038/s41522-025-00903-7

Aitchison, J. (1982). The Statistical Analysis of Compositional Data. Journal of the Royal Statistical Society. Series B (Methodological), 44(2), 139–177.

Aitchison, J., Barceló-Vidal, C., Martín-Fernández, J. A., & Pawlowsky-Glahn, V. (2000). Logratio Analysis and Compositional Distance. Mathematical Geology, 32(3), 271–275. 10.1023/A:1007529726302

Amir, A., McDonald, D., Navas-Molina, J. A., Kopylova, E., Morton, J. T., Zech Xu, Z., Kightley, E. P., Thompson, L. R., Hyde, E. R., Gonzalez, A., & Knight, R. (2017). Deblur Rapidly Resolves Single-Nucleotide Community Sequence Patterns. mSystems, 2(2). 10.1128/mSystems.00191-16

Andrews, S. (2010). FastQC: A Quality Control tool for High Throughput Sequence Data. https://www.bioinformatics.babraham.ac.uk/projects/fastqc/

Ankenbauer, K. E., Yang, Y., Chung, C.-Y., Andrade, L. R., Novak, S. W., Jarvis, B., Ali Hanel, W. H., Liu, J., Sarkisian, V., Dani, N., Krystofiak, E., Hu, G., Ebrahim, S., Kachar, B., Gong, Q., Wahl, G. M., Lau, K. S., Brown, J. W., Manor, U., & DelGiorno, K. E. (2026). Protocadherin 20 Is a POU Class 2 Homeobox 3 Target Gene Required for Proper Tuft Cell Microvillus Organization. Cellular and Molecular Gastroenterology and Hepatology, 20(6), 101742. 10.1016/j.jcmgh.2026.101742

Aoyama, K., Yamamura, R., Katsurada, T., Shimizu, T., Takahashi, D., Kondo, E., Iwasaki, N., Tamakoshi, A., Soga, T., Fukuda, S., Sonoshita, M., & Sakamoto, N. (2025). Decreased Fecal Nicotinamide and Increased Bacterial Nicotinamidase Gene Expression in Ulcerative Colitis Patients. *Inflammatory Bowel Diseases*, izaf092. 10.1093/ibd/izaf092

Bates, D., Maechler, M., Bolker [aut, B., cre, Walker, S., Christensen, R. H. B., Singmann, H., Dai, B., Scheipl, F., Grothendieck, G., Green, P., Fox, J., Bauer, A., simulate.formula), P. N. K. (shared copyright on, Tanaka, E., Jagan, M., & Boylan, R. D. (2025). lme4: Linear Mixed-Effects Models using “Eigen” and S4 (Version 1.1-37) [Computer software]. https://cran.r-project.org/web/packages/lme4/index.html

Bates, M. D., Erwin, C. R., Sanford, L. P., Wiginton, D., Bezerra, J. A., Schatzman, L. C., Jegga, A. G., Ley-Ebert, C., Williams, S. S., Steinbrecher, K. A., Warner, B. W., Cohen, M. B., & Aronow, B. J. (2002). Novel genes and functional relationships in the adult mouse gastrointestinal tract identified by microarray analysis. Gastroenterology, 122(5), 1467–1482. 10.1053/gast.2002.32975

Benjamini, Y., & Hochberg, Y. (1995). Controlling the False Discovery Rate: A Practical and Powerful Approach to Multiple Testing. Journal of the Royal Statistical Society. Series B (Methodological), 57(1), 289–300.

Bhat, A. A., Nisar, S., Singh, M., Ashraf, B., Masoodi, T., Prasad, C. P., Sharma, A., Maacha, S., Karedath, T., Hashem, S., Yasin, S. B., Bagga, P., Reddy, R., Frennaux, M. P., Uddin, S., Dhawan, P., Haris, M., & Macha, M. A. (2022). Cytokine- and chemokine-induced inflammatory colorectal tumor microenvironment: Emerging avenue for targeted therapy. Cancer Communications, 42(8), 689–715. 10.1002/cac2.12295

Bisanz, J. E. (2018). qiime2R: Importing QIIME2 artifacts and associated data into R sessions. (Version v0.99) [Computer software]. https://github.com/jbisanz/qiime2R

Blekhman, R., Goodrich, J. K., Huang, K., Sun, Q., Bukowski, R., Bell, J. T., Spector, T. D., Keinan, A., Ley, R. E., Gevers, D., & Clark, A. G. (2015). Host genetic variation impacts microbiome composition across human body sites. Genome Biology, 16(1), 191. 10.1186/s13059-015-0759-1

Blesl, A., Binder, L., Halwachs, B., Baumann-Durchschein, F., Fürst, S., Constantini-Kump, P., Wenzl, H., Gorkiewicz, G., & Högenauer, C. (2025). The Fecal Microbiome of IBD Patients Is Less Divertible by Bowel Preparation Compared to Healthy Controls: Results From a Prospective Study. Inflammatory Bowel Diseases, 31(7), 2007–2018. 10.1093/ibd/izaf053

Bokulich, N. A., Kaehler, B. D., Rideout, J. R., Dillon, M., Bolyen, E., Knight, R., Huttley, G. A., & Gregory Caporaso, J. (2018). Optimizing taxonomic classification of marker-gene amplicon sequences with QIIME 2’s q2-feature-classifier plugin. Microbiome, 6(1), 90. 10.1186/s40168-018-0470-z

Bokulich, N. A., Subramanian, S., Faith, J. J., Gevers, D., Gordon, J. I., Knight, R., Mills, D. A., & Caporaso, J. G. (2013). Quality-filtering vastly improves diversity estimates from Illumina amplicon sequencing. Nature Methods, 10(1), 57–59. 10.1038/nmeth.2276

Bolyen, E., Rideout, J. R., Dillon, M. R., Bokulich, N. A., Abnet, C. C., Al-Ghalith, G. A., Alexander, H., Alm, E. J., Arumugam, M., Asnicar, F., Bai, Y., Bisanz, J. E., Bittinger, K., Brejnrod, A., Brislawn, C. J., Brown, C. T., Callahan, B. J., Caraballo-Rodríguez, A. M., Chase, J., … Caporaso, J. G. (2019). Reproducible, interactive, scalable and extensible microbiome data science using QIIME 2. Nature Biotechnology, 37(8), 852–857. 10.1038/s41587-019-0209-9

Bonder, M. J., Kurilshikov, A., Tigchelaar, E. F., Mujagic, Z., Imhann, F., Vila, A. V., Deelen, P., Vatanen, T., Schirmer, M., Smeekens, S. P., Zhernakova, D. V., Jankipersadsing, S. A., Jaeger, M., Oosting, M., Cenit, M. C., Masclee, A. A. M., Swertz, M. A., Li, Y., Kumar, V., … Zhernakova, A. (2016). The effect of host genetics on the gut microbiome. Nature Genetics, 48(11), Article 11. 10.1038/ng.3663

Boni, C., Laudanna, C., & Sorio, C. (2022). A Comprehensive Review of Receptor-Type Tyrosine-Protein Phosphatase Gamma (PTPRG) Role in Health and Non-Neoplastic Disease. Biomolecules, 12(1), 84. 10.3390/biom12010084

Boyle, E. I., Weng, S., Gollub, J., Jin, H., Botstein, D., Cherry, J. M., & Sherlock, G. (2004). GO::TermFinder—Open source software for accessing Gene Ontology information and finding significantly enriched Gene Ontology terms associated with a list of genes. Bioinformatics, 20(18), 3710–3715. 10.1093/bioinformatics/bth456

Bray, J. R., & Curtis, J. T. (1957). An Ordination of the Upland Forest Communities of Southern Wisconsin. Ecological Monographs, 27(4), 325–349. 10.2307/1942268

Bubier, J. A., Chesler, E. J., & Weinstock, G. M. (2021). Host genetic control of gut microbiome composition. Mammalian Genome: Official Journal of the International Mammalian Genome Society, 32(4), 263–281. 10.1007/s00335-021-09884-2

Burclaff, J., Bliton, R. J., Breau, K. A., Ok, M. T., Gomez-Martinez, I., Ranek, J. S., Bhatt, A. P., Purvis, J. E., Woosley, J. T., & Magness, S. T. (2022). A Proximal-to-Distal Survey of Healthy Adult Human Small Intestine and Colon Epithelium by Single-Cell Transcriptomics. Cellular and Molecular Gastroenterology and Hepatology, 13(5), 1554–1589. 10.1016/j.jcmgh.2022.02.007

Bush, S. J., Connor, T. R., Peto, T. E. A., Crook, D. W., & Walker, A. S. (2020). Evaluation of methods for detecting human reads in microbial sequencing datasets. Microbial Genomics, 6(7), mgen000393. 10.1099/mgen.0.000393

Cabrera-Rivera, G. L., Madera-Sandoval, R. L., León-Pedroza, J. I., Ferat-Osorio, E., Salazar-Rios, E., Hernández-Aceves, J. A., Guadarrama-Aranda, U., López-Macías, C., Wong-Baeza, I., & Arriaga-Pizano, L. A. (2022). Increased TNF-α production in response to IL-6 in patients with systemic inflammation without infection. Clinical and Experimental Immunology, 209(2), 225–235. 10.1093/cei/uxac055

Cai, C., Zhu, S., Tong, J., Wang, T., Feng, Q., Qiao, Y., & Shen, J. (2021). Relating the transcriptome and microbiome by paired terminal ileal Crohn disease. iScience, 24(6), 102516. 10.1016/j.isci.2021.102516

Caporaso, J. G., Lauber, C. L., Walters, W. A., Berg-Lyons, D., Huntley, J., Fierer, N., Owens, S. M., Betley, J., Fraser, L., Bauer, M., Gormley, N., Gilbert, J. A., Smith, G., & Knight, R. (2012). Ultra-high-throughput microbial community analysis on the Illumina HiSeq and MiSeq platforms. The ISME Journal, 6(8), Article 8. 10.1038/ismej.2012.8

Carroll, I. M., Chang, Y.-H., Park, J., Sartor, R. B., & Ringel, Y. (2010). Luminal and mucosal-associated intestinal microbiota in patients with diarrhea-predominant irritable bowel syndrome. Gut Pathogens, 2(1), 19. 10.1186/1757-4749-2-19

Chavez-Arroyo, A., Radlinski, L. C., & Bäumler, A. J. (2025). Principles of gut microbiota assembly. Trends in Microbiology, 33(7), 718–726. 10.1016/j.tim.2025.02.014

Chen, W., Liu, F., Ling, Z., Tong, X., & Xiang, C. (2012). Human Intestinal Lumen and Mucosa-Associated Microbiota in Patients with Colorectal Cancer. PLOS ONE, 7(6), e39743. 10.1371/journal.pone.0039743

Comelli, E. M., Lariani, S., Zwahlen, M.-C., Fotopoulos, G., Holzwarth, J. A., Cherbut, C., Dorta, G., Corthésy-Theulaz, I., & Grigorov, M. (2009). Biomarkers of human gastrointestinal tract regions. Mammalian Genome: Official Journal of the International Mammalian Genome Society, 20(8), 516–527. 10.1007/s00335-009-9212-7

Dave, M., Johnson, L. A., Walk, S. T., Young, V. B., Stidham, R. W., Chaudhary, M. N., FunNell, J., & Higgins, P. D. R. (2011). A randomised trial of sheathed versus standard forceps for obtaining uncontaminated biopsy specimens of microbiota from the terminal ileum. Gut, 60(8), 1043–1049. 10.1136/gut.2010.224337

Davenport, E. R., Cusanovich, D. A., Michelini, K., Barreiro, L. B., Ober, C., & Gilad, Y. (2015). Genome-Wide Association Studies of the Human Gut Microbiota. PLOS ONE, 10(11), e0140301. 10.1371/journal.pone.0140301

Davis, N. M., Proctor, D. M., Holmes, S. P., Relman, D. A., & Callahan, B. J. (2018). Simple statistical identification and removal of contaminant sequences in marker-gene and metagenomics data. Microbiome, 6(1), 226. 10.1186/s40168-018-0605-2

De Maesschalck, C., Van Immerseel, F., Eeckhaut, V., De Baere, S., Cnockaert, M., Croubels, S., Haesebrouck, F., Ducatelle, R., & Vandamme, P. (2014). Faecalicoccus acidiformans gen. Nov., sp. Nov., isolated from the chicken caecum, and reclassification of Streptococcus pleomorphus (Barnes et al. 1977), Eubacterium biforme (Eggerth 1935) and Eubacterium cylindroides (Cato et al. 1974) as Faecalicoccus pleomorphus comb. Nov., Holdemanella biformis gen. Nov., comb. Nov. And Faecalitalea cylindroides gen. Nov., comb. Nov., respectively, within the family Erysipelotrichaceae. International Journal of Systematic and Evolutionary Microbiology, 64(Pt_11), 3877–3884. 10.1099/ijs.0.064626-0

Dixon, P. (2003). VEGAN, a package of R functions for community ecology. Journal of Vegetation Science, 14(6), 927–930. 10.1111/j.1654-1103.2003.tb02228.x

Dobin, A., Davis, C. A., Schlesinger, F., Drenkow, J., Zaleski, C., Jha, S., Batut, P., Chaisson, M., & Gingeras, T. R. (2013). STAR: Ultrafast universal RNA-seq aligner. Bioinformatics, 29(1), 15–21. 10.1093/bioinformatics/bts635

Dworkin, M., Falkow, S., Rosenberg, E., Schleifer, K.-H., & Stackebrandt, E. (Eds.). (2006). The Prokaryotes: Volume 4: Bacteria: Firmicutes, Cyanobacteria. Springer US. 10.1007/0-387-30744-3

Englander, E. W. (2005). Gene expression changes reveal patterns of aging in the rat digestive tract. *Ageing Research Reviews*, Gene Expression During Aging, 4(4), 564–578. 10.1016/j.arr.2005.06.005

Faith, D. P. (1992). Conservation evaluation and phylogenetic diversity. Biological Conservation, 61(1), 1–10. 10.1016/0006-3207(92)91201-3

Fan, Y., & Pedersen, O. (2021). Gut microbiota in human metabolic health and disease. Nature Reviews Microbiology, 19(1), 55–71. 10.1038/s41579-020-0433-9

Ferretti, P., Johnson, K., Priya, S., & Blekhman, R. (2025). Genomics of host–microbiome interactions in humans. Nature Reviews Genetics, 1–19. 10.1038/s41576-025-00849-8

Ferrie, S., Webster, A., Wu, B., Tan, C., & Carey, S. (2021). Gastrointestinal surgery and the gut microbiome: A systematic literature review. European Journal of Clinical Nutrition, 75(1), 12–25. 10.1038/s41430-020-0681-9

Fierer, N., Leung, P. M., Lappan, R., Eisenhofer, R., Ricci, F., Holland, S. I., Dragone, N., Blackall, L. L., Dong, X., Dorador, C., Ferrari, B. C., Goordial, J., Holmes, S. P., Inagaki, F., Korem, T., Li, S. S., Makhalanyane, T. P., Metcalf, J. L., Nagarajan, N., … Greening, C. (2025). Guidelines for preventing and reporting contamination in low-biomass microbiome studies. Nature Microbiology, 10(7), 1570–1580. 10.1038/s41564-025-02035-2

Fink, M., & Wrana, J. L. (2023). Regulation of homeostasis and regeneration in the adult intestinal epithelium by the TGF-β superfamily. Developmental Dynamics, 252(4), 445–462. 10.1002/dvdy.500

Fisher, R. A., Corbet, A. S., & Williams, C. B. (1943). The Relation Between the Number of Species and the Number of Individuals in a Random Sample of an Animal Population. Journal of Animal Ecology, 12(1), 42–58. 10.2307/1411

Folashade, D., Ooi, H., Calaway, R., & Weston, S. (2022). foreach: Provides Foreach Looping Construct (Version 1.5.2) [Computer software]. https://cran.r-project.org/web/packages/foreach/index.html

Folashade, D., Weston, S., & Tenenbaum, D. (2022). doParallel: Foreach Parallel Adaptor for the “parallel” Package (Version 1.0.17) [Computer software]. https://cran.r-project.org/web/packages/doParallel/index.html

Friedman, J. H., Hastie, T., & Tibshirani, R. (2010). Regularization Paths for Generalized Linear Models via Coordinate Descent. Journal of Statistical Software, 33, 1–22. 10.18637/jss.v033.i01

Fujimiya, M., & Inui, A. (2000). Peptidergic regulation of gastrointestinal motility in rodents. Peptides, 21(10), 1565–1582. 10.1016/S0196-9781(00)00313-2

Fung, C., Fraser, L. M., Barrón, G. M., Gologorsky, M. B., Atkinson, S. N., Gerrick, E. R., Hayward, M., Ziegelbauer, J., Li, J. A., Nico, K. F., Tyner, M. D. W., DeSchepper, L. B., Pan, A., Salzman, N. H., & Howitt, M. R. (2023). Tuft cells mediate commensal remodeling of the small intestinal antimicrobial landscape. Proceedings of the National Academy of Sciences of the United States of America, 120(23), e2216908120. 10.1073/pnas.2216908120

Gardiner, B. J., Tai, A. Y., Kotsanas, D., Francis, M. J., Roberts, S. A., Ballard, S. A., Junckerstorff, R. K., & Korman, T. M. (2015). Clinical and Microbiological Characteristics of Eggerthella lenta Bacteremia. Journal of Clinical Microbiology, 53(2), 626–635. 10.1128/JCM.02926-14

Ghaffari, S., Abbasi, A., Somi, M. H., Moaddab, S. Y., Nikniaz, L., Kafil, H. S., & Ebrahimzadeh Leylabadlo, H. (2022). Akkermansia muciniphila: From its critical role in human health to strategies for promoting its abundance in human gut microbiome. Critical Reviews in Food Science and Nutrition, 0(0), 1–21. 10.1080/10408398.2022.2045894

Glebov, O. K., Rodriguez, L. M., Nakahara, K., Jenkins, J., Cliatt, J., Humbyrd, C.-J., DeNobile, J., Soballe, P., Simon, R., Wright, G., Lynch, P., Patterson, S., Lynch, H., Gallinger, S., Buchbinder, A., Gordon, G., Hawk, E., & Kirsch, I. R. (2003). Distinguishing right from left colon by the pattern of gene expression. *Cancer Epidemiology, Biomarkers & Prevention: A Publication of the American Association for Cancer Research*, Cosponsored by the American Society of Preventive Oncology, 12(8), 755–762.

Goodrich, J. K., Davenport, E. R., Beaumont, M., Jackson, M. A., Knight, R., Ober, C., Spector, T. D., Bell, J. T., Clark, A. G., & Ley, R. E. (2016). Genetic Determinants of the Gut Microbiome in UK Twins. Cell Host & Microbe, 19(5), 731–743. 10.1016/j.chom.2016.04.017

Goudarzi, A. (2019). The recent insights into the function of ACAT1: A possible anti-cancer therapeutic target. Life Sciences, 232, 116592. 10.1016/j.lfs.2019.116592

Gower, J. C. (1975). Generalized procrustes analysis. Psychometrika, 40(1), 33–51. 10.1007/BF02291478

Hickey, J. W., Becker, W. R., Nevins, S. A., Horning, A., Perez, A. E., Zhu, C., Zhu, B., Wei, B., Chiu, R., Chen, D. C., Cotter, D. L., Esplin, E. D., Weimer, A. K., Caraccio, C., Venkataraaman, V., Schürch, C. M., Black, S., Brbić, M., Cao, K., … Snyder, M. (2023). Organization of the human intestine at single-cell resolution. Nature, 619(7970), 572–584. 10.1038/s41586-023-05915-x

Hidalgo-Cantabrana, C., Delgado, S., Ruiz, L., Ruas-Madiedo, P., Sánchez, B., & Margolles, A. (2017). Bifidobacteria and Their Health-Promoting Effects. Microbiology Spectrum, 5(3). 10.1128/microbiolspec.BAD-0010-2016

Hoffman, G. E., & Roussos, P. (2020). Dream: Powerful differential expression analysis for repeated measures designs. Bioinformatics, 37(2), 192–201. 10.1093/bioinformatics/btaa687

Hoffman, G. E., & Schadt, E. E. (2016). variancePartition: Interpreting drivers of variation in complex gene expression studies. BMC Bioinformatics, 17(1), 483. 10.1186/s12859-016-1323-z

Hu, S., Bourgonje, A. R., Gacesa, R., Jansen, B. H., Björk, J. R., Bangma, A., Hidding, I. J., van Dullemen, H. M., Visschedijk, M. C., Faber, K. N., Dijkstra, G., Harmsen, H. J. M., Festen, E. A. M., Vich Vila, A., Spekhorst, L. M., & Weersma, R. K. (2024). Mucosal host-microbe interactions associate with clinical phenotypes in inflammatory bowel disease. Nature Communications, 15(1), 1470. 10.1038/s41467-024-45855-2

Huang, S., Xie, Z., Han, J., Wang, H., Yang, G., Li, M., Zhou, G., Wang, Y., Li, L., Li, L., Zeng, Z., Yu, J., Chen, M., & Zhang, S. (2023). Protocadherin 20 maintains intestinal barrier function to protect against Crohn’s disease by targeting ATF6. Genome Biology, 24(1), 159. 10.1186/s13059-023-02991-0

Ishida, S., Kato, K., Tanaka, M., Odamaki, T., Kubo, R., Mitsuyama, E., Xiao, J., Yamaguchi, R., Uematsu, S., Imoto, S., & Miyano, S. (2020). Genome-wide association studies and heritability analysis reveal the involvement of host genetics in the Japanese gut microbiota. Communications Biology, 3(1), 686. 10.1038/s42003-020-01416-z

James, K. R., Gomes, T., Elmentaite, R., Kumar, N., Gulliver, E. L., King, H. W., Stares, M. D., Bareham, B. R., Ferdinand, J. R., Petrova, V. N., Polański, K., Forster, S. C., Jarvis, L. B., Suchanek, O., Howlett, S., James, L. K., Jones, J. L., Meyer, K. B., Clatworthy, M. R., … Teichmann, S. A. (2020). Distinct microbial and immune niches of the human colon. Nature Immunology, 21(3), Article 3. 10.1038/s41590-020-0602-z

Jangi, S., Gandhi, R., Cox, L. M., Li, N., von Glehn, F., Yan, R., Patel, B., Mazzola, M. A., Liu, S., Glanz, B. L., Cook, S., Tankou, S., Stuart, F., Melo, K., Nejad, P., Smith, K., Topçuolu, B. D., Holden, J., Kivisäkk, P., … Weiner, H. L. (2016). Alterations of the human gut microbiome in multiple sclerosis. Nature Communications, 7(1), Article 1. 10.1038/ncomms12015

Jensen, B. A. H., Heyndrickx, M., Jonkers, D., Mackie, A., Millet, S., Naghibi, M., Pærregaard, S. I., Pot, B., Saulnier, D., Sina, C., Sterkman, L. G. W., Van den Abbeele, P., Venlet, N. V., Zoetendal, E. G., & Ouwehand, A. C. (2023). Small intestine vs. colon ecology and physiology: Why it matters in probiotic administration. Cell Reports Medicine, 4(9), 101190. 10.1016/j.xcrm.2023.101190

Jin, M., Kalainy, S., Baskota, N., Chiang, D., Deehan, E. C., McDougall, C., Tandon, P., Martínez, I., Cervera, C., Walter, J., & Abraldes, J. G. (2019). Faecal microbiota from patients with cirrhosis has a low capacity to ferment non-digestible carbohydrates into short-chain fatty acids. Liver International, 39(8), 1437–1447. 10.1111/liv.14106

Juge, N. (2022). Relationship between mucosa-associated gut microbiota and human diseases. Biochemical Society Transactions, 50(5), 1225–1236. 10.1042/BST20201201

Kanehisa, M., & Goto, S. (2000). KEGG: Kyoto encyclopedia of genes and genomes. Nucleic Acids Research, 28(1), 27–30. 10.1093/nar/28.1.27

Kastl, A. J., Terry, N. A., Wu, G. D., & Albenberg, L. G. (2019). The Structure and Function of the Human Small Intestinal Microbiota: Current Understanding and Future Directions. Cellular and Molecular Gastroenterology and Hepatology, 9(1), 33–45. 10.1016/j.jcmgh.2019.07.006

Katoh, K., Misawa, K., Kuma, K., & Miyata, T. (2002). MAFFT: A novel method for rapid multiple sequence alignment based on fast Fourier transform. Nucleic Acids Research, 30(14), 3059–3066. 10.1093/nar/gkf436

Kim, N., Gim, J.-A., Lee, B. J., Choi, B. il, Yoon, H. S., Kim, S. H., Joo, M. K., Park, J.-J., & Kim, C. (2022). Crosstalk between mucosal microbiota, host gene expression, and sociomedical factors in the progression of colorectal cancer. Scientific Reports, 12(1), 13447. 10.1038/s41598-022-17823-7

Kuhn, M. (2008). Building Predictive Models in R Using the caret Package. Journal of Statistical Software, 28, 1–26. 10.18637/jss.v028.i05

Kurilshikov, A., Medina-Gomez, C., Bacigalupe, R., Radjabzadeh, D., Wang, J., Demirkan, A., Le Roy, C. I., Raygoza Garay, J. A., Finnicum, C. T., Liu, X., Zhernakova, D. V., Bonder, M. J., Hansen, T. H., Frost, F., Rühlemann, M. C., Turpin, W., Moon, J.-Y., Kim, H.-N., Lüll, K., … Zhernakova, A. (2021). Large-scale association analyses identify host factors influencing human gut microbiome composition. Nature Genetics, 53(2), Article 2. 10.1038/s41588-020-00763-1

Lahti, L., & Shetty, S. (2017). Microbiome R package. 10.18129/B9.bioc.microbiome

Langmead, B., & Salzberg, S. L. (2012). Fast gapped-read alignment with Bowtie 2. Nature Methods, 9(4), 357–359. 10.1038/nmeth.1923

Leek, J. T., Scharpf, R. B., Bravo, H. C., Simcha, D., Langmead, B., Johnson, W. E., Geman, D., Baggerly, K., & Irizarry, R. A. (2010). Tackling the widespread and critical impact of batch effects in high-throughput data. Nature Reviews Genetics, 11(10), 733–739. 10.1038/nrg2825

Leite, G. G. S., Weitsman, S., Parodi, G., Celly, S., Sedighi, R., Sanchez, M., Morales, W., Villanueva-Millan, M. J., Barlow, G. M., Mathur, R., Lo, S. K., Jamil, L. H., Paski, S., Rezaie, A., & Pimentel, M. (2020). Mapping the Segmental Microbiomes in the Human Small Bowel in Comparison with Stool: A REIMAGINE Study. Digestive Diseases and Sciences, 65(9), 2595–2604. 10.1007/s10620-020-06173-x

Li, G., Yang, M., Zhou, K., Zhang, L., Tian, L., Lv, S., Jin, Y., Qian, W., Xiong, H., Lin, R., Fu, Y., & Hou, X. (2015). Diversity of Duodenal and Rectal Microbiota in Biopsy Tissues and Luminal Contents in Healthy Volunteers. 25(7), 1136–1145. 10.4014/jmb.1412.12047

Liao, Y., Smyth, G. K., & Shi, W. (2014). featureCounts: An efficient general purpose program for assigning sequence reads to genomic features. Bioinformatics, 30(7), 923–930. 10.1093/bioinformatics/btt656

Lim, M. Y., You, H. J., Yoon, H. S., Kwon, B., Lee, J. Y., Lee, S., Song, Y.-M., Lee, K., Sung, J., & Ko, G. (2017). The effect of heritability and host genetics on the gut microbiota and metabolic syndrome. Gut, 66(6), 1031–1038. 10.1136/gutjnl-2015-311326

Liu, P., Wang, Y., Yang, G., Zhang, Q., Meng, L., Xin, Y., & Jiang, X. (2021). The role of short-chain fatty acids in intestinal barrier function, inflammation, oxidative stress, and colonic carcinogenesis. Pharmacological Research, 165, 105420. 10.1016/j.phrs.2021.105420

Liu, T., Zhang, L., Joo, D., & Sun, S.-C. (2017). NF-κB signaling in inflammation. Signal Transduction and Targeted Therapy, 2(1), 17023. 10.1038/sigtrans.2017.23

Lkhagva, E., Chung, H.-J., Hong, J., Tang, W. H. W., Lee, S.-I., Hong, S.-T., & Lee, S. (2021). The regional diversity of gut microbiome along the GI tract of male C57BL/6 mice. BMC Microbiology, 21(1), 44. 10.1186/s12866-021-02099-0

Lloyd-Price, J., Arze, C., Ananthakrishnan, A. N., Schirmer, M., Avila-Pacheco, J., Poon, T. W., Andrews, E., Ajami, N. J., Bonham, K. S., Brislawn, C. J., Casero, D., Courtney, H., Gonzalez, A., Graeber, T. G., Hall, A. B., Lake, K., Landers, C. J., Mallick, H., Plichta, D. R., … Huttenhower, C. (2019). Multi-omics of the gut microbial ecosystem in inflammatory bowel diseases. Nature, 569(7758), Article 7758. 10.1038/s41586-019-1237-9

Lopera-Maya, E. A., Kurilshikov, A., van der Graaf, A., Hu, S., Andreu-Sánchez, S., Chen, L., Vila, A. V., Gacesa, R., Sinha, T., Collij, V., Klaassen, M. A. Y., Bolte, L. A., Gois, M. F. B., Neerincx, P. B. T., Swertz, M. A., Harmsen, H. J. M., Wijmenga, C., Fu, J., Weersma, R. K., … Sanna, S. (2022). Effect of host genetics on the gut microbiome in 7,738 participants of the Dutch Microbiome Project. Nature Genetics, 54(2), Article 2. 10.1038/s41588-021-00992-y

Love, M. I., Huber, W., & Anders, S. (2014). Moderated estimation of fold change and dispersion for RNA-seq data with DESeq2. Genome Biology, 15(12), 550. 10.1186/s13059-014-0550-8

Lozupone, C., & Knight, R. (2005). UniFrac: A New Phylogenetic Method for Comparing Microbial Communities. Applied and Environmental Microbiology, 71(12), 8228–8235. 10.1128/AEM.71.12.8228-8235.2005

Ma, Q., Li, Y., Wang, J., Li, P., Duan, Y., Dai, H., An, Y., Cheng, L., Wang, T., Wang, C., Wang, T., & Zhao, B. (2020). Investigation of gut microbiome changes in type 1 diabetic mellitus rats based on high-throughput sequencing. Biomedicine & Pharmacotherapy, 124, 109873. 10.1016/j.biopha.2020.109873

Mallick, H., Rahnavard, A., McIver, L. J., Ma, S., Zhang, Y., Nguyen, L. H., Tickle, T. L., Weingart, G., Ren, B., Schwager, E. H., Chatterjee, S., Thompson, K. N., Wilkinson, J. E., Subramanian, A., Lu, Y., Waldron, L., Paulson, J. N., Franzosa, E. A., Bravo, H. C., & Huttenhower, C. (2021). Multivariable association discovery in population-scale meta-omics studies. PLoS Computational Biology, 17(11), e1009442. 10.1371/journal.pcbi.1009442

Mankarious, M. M., Connelly, T. M., Harris, L., Deiling, S., Yochum, G. S., & Koltun, W. A. (2023). Creating a Surgical Biobank: The Hershey Medical Center Experience. Diseases of the Colon and Rectum, 66(9), 1174–1184. 10.1097/DCR.0000000000002944

Martin, M. (2011). Cutadapt removes adapter sequences from high-throughput sequencing reads. EMBnet.Journal, 17(1), Article 1. 10.14806/ej.17.1.200

Martinez-Guryn, K., Leone, V., & Chang, E. B. (2019). Regional Diversity of the Gastrointestinal Microbiome. Cell Host & Microbe, 26(3), 314–324. 10.1016/j.chom.2019.08.011

McMurdie, P. J., & Holmes, S. (2013). phyloseq: An R Package for Reproducible Interactive Analysis and Graphics of Microbiome Census Data. PLOS ONE, 8(4), e61217. 10.1371/journal.pone.0061217

Medawar, E., Haange, S.-B., Rolle-Kampczyk, U., Engelmann, B., Dietrich, A., Thieleking, R., Wiegank, C., Fries, C., Horstmann, A., Villringer, A., von Bergen, M., Fenske, W., & Veronica Witte, A. (2021). Gut microbiota link dietary fiber intake and short-chain fatty acid metabolism with eating behavior. Translational Psychiatry, 11(1), 500. 10.1038/s41398-021-01620-3

Mölder, F., Jablonski, K. P., Letcher, B., Hall, M. B., Tomkins-Tinch, C. H., Sochat, V., Forster, J., Lee, S., Twardziok, S. O., Kanitz, A., Wilm, A., Holtgrewe, M., Rahmann, S., Nahnsen, S., & Köster, J. (2021). *Sustainable data analysis with Snakemake* (10:33). F1000Research. 10.12688/f1000research.29032.2

Mowat, A. M., & Agace, W. W. (2014). Regional specialization within the intestinal immune system. Nature Reviews Immunology, 14(10), 667–685. 10.1038/nri3738

Nichols, R. G., & Davenport, E. R. (2021). The relationship between the gut microbiome and host gene expression: A review. Human Genetics, 140(5), 747–760. 10.1007/s00439-020-02237-0

O’Brien, C. L., Allison, G. E., Grimpen, F., & Pavli, P. (2013). Impact of Colonoscopy Bowel Preparation on Intestinal Microbiota. PLOS ONE, 8(5), e62815. 10.1371/journal.pone.0062815

Oh, D. S., Lieu, S. N., Yamaguchi, D. J., Tachiki, K., Lambrecht, N., Ohning, G. V., Sachs, G., Germano, P. M., & Pisegna, J. R. (2005). PACAP regulation of secretion and proliferation of pure populations of gastric ECL cells. Journal of Molecular Neuroscience, 26(1), 85–98. 10.1385/JMN:26:1:085

Oliphant, K., & Allen-Vercoe, E. (2019). Macronutrient metabolism by the human gut microbiome: Major fermentation by-products and their impact on host health. Microbiome, 7(1), 91. 10.1186/s40168-019-0704-8

Onyeaka, H. N., & Nwabor, O. F. (2022). Lactic acid bacteria and bacteriocins as biopreservatives. In Food Preservation and Safety of Natural Products (pp. 147–162). Elsevier. 10.1016/B978-0-323-85700-0.00012-5

Parada Venegas, D., De la Fuente, M. K., Landskron, G., González, M. J., Quera, R., Dijkstra, G., Harmsen, H. J. M., Faber, K. N., & Hermoso, M. A. (2019). Short Chain Fatty Acids (SCFAs)-Mediated Gut Epithelial and Immune Regulation and Its Relevance for Inflammatory Bowel Diseases. Frontiers in Immunology, 10. 10.3389/fimmu.2019.00277

Pérez-Reytor, D., Puebla, C., Karahanian, E., & García, K. (2021). Use of Short-Chain Fatty Acids for the Recovery of the Intestinal Epithelial Barrier Affected by Bacterial Toxins. Frontiers in Physiology, 12, 650313. 10.3389/fphys.2021.650313

Pihelgas, S., Ehala-Aleksejev, K., Adamberg, S., Kazantseva, J., & Adamberg, K. (2024). The gut microbiota of healthy individuals remains resilient in response to the consumption of various dietary fibers. Scientific Reports, 14(1), 22208. 10.1038/s41598-024-72673-9

Price, M. N., Dehal, P. S., & Arkin, A. P. (2010). FastTree 2 – Approximately Maximum-Likelihood Trees for Large Alignments. PLOS ONE, 5(3), e9490. 10.1371/journal.pone.0009490

Priya, S., Burns, M. B., Ward, T., Mars, R. A. T., Adamowicz, B., Lock, E. F., Kashyap, P. C., Knights, D., & Blekhman, R. (2022). Identification of shared and disease-specific host gene–microbiome associations across human diseases using multi-omic integration. Nature Microbiology, 7(6), 780–795. 10.1038/s41564-022-01121-z

Qin, Y., Havulinna, A. S., Liu, Y., Jousilahti, P., Ritchie, S. C., Tokolyi, A., Sanders, J. G., Valsta, L., Brożyńska, M., Zhu, Q., Tripathi, A., Vázquez-Baeza, Y., Loomba, R., Cheng, S., Jain, M., Niiranen, T., Lahti, L., Knight, R., Salomaa, V., … Méric, G. (2022). Combined effects of host genetics and diet on human gut microbiota and incident disease in a single population cohort. Nature Genetics, 54(2), Article 2. 10.1038/s41588-021-00991-z

Quast, C., Pruesse, E., Yilmaz, P., Gerken, J., Schweer, T., Yarza, P., Peplies, J., & Glöckner, F. O. (2013). The SILVA ribosomal RNA gene database project: Improved data processing and web-based tools. Nucleic Acids Research, 41(D1), D590–D596. 10.1093/nar/gks1219

Rangel, I., Sundin, J., Fuentes, S., Repsilber, D., de Vos, W. M., & Brummer, R. J. (2015). The relationship between faecal-associated and mucosal-associated microbiota in irritable bowel syndrome patients and healthy subjects. Alimentary Pharmacology & Therapeutics, 42(10), 1211–1221. 10.1111/apt.13399

Robinson, M. D., McCarthy, D. J., & Smyth, G. K. (2010). edgeR: A Bioconductor package for differential expression analysis of digital gene expression data. Bioinformatics, 26(1), 139–140. 10.1093/bioinformatics/btp616

Rognes, T., Flouri, T., Nichols, B., Quince, C., & Mahé, F. (2016). VSEARCH: A versatile open source tool for metagenomics. PeerJ, 4, e2584. 10.7717/peerj.2584

Rowland, I., Gibson, G., Heinken, A., Scott, K., Swann, J., Thiele, I., & Tuohy, K. (2018). Gut microbiota functions: Metabolism of nutrients and other food components. European Journal of Nutrition, 57(1), 1–24. 10.1007/s00394-017-1445-8

Ruigrok, R. A. A. A., Weersma Rinse K., & and Vich Vila, A. (2023). The emerging role of the small intestinal microbiota in human health and disease. Gut Microbes, 15(1), 2201155. 10.1080/19490976.2023.2201155

Schaefer, C. F., Anthony, K., Krupa, S., Buchoff, J., Day, M., Hannay, T., & Buetow, K. H. (2009). PID: The Pathway Interaction Database. Nucleic Acids Research, 37(Database issue), D674–D679. 10.1093/nar/gkn653

Scott, S. A., Fu, J., & Chang, P. V. (2020). Microbial tryptophan metabolites regulate gut barrier function via the aryl hydrocarbon receptor. Proceedings of the National Academy of Sciences, 117(32), 19376–19387. 10.1073/pnas.2000047117

Shalon, D., Culver, R. N., Grembi, J. A., Folz, J., Treit, P. V., Shi, H., Rosenberger, F. A., Dethlefsen, L., Meng, X., Yaffe, E., Aranda-Díaz, A., Geyer, P. E., Mueller-Reif, J. B., Spencer, S., Patterson, A. D., Triadafilopoulos, G., Holmes, S. P., Mann, M., Fiehn, O., … Huang, K. C. (2023). Profiling the human intestinal environment under physiological conditions. Nature, 617(7961), 581–591. 10.1038/s41586-023-05989-7

Shannon, C. E. (1948). A Mathematical Theory of Communication. Bell System Technical Journal, 27(3), 379–423. 10.1002/j.1538-7305.1948.tb01338.x

Shin, Y.-H., Bang, S., Xavier, R., & Clardy, J. (2025). Eggerthella lenta Produces a Cryptic Pro-inflammatory Lipid. Journal of the American Chemical Society, 147(29), 25180–25183. 10.1021/jacs.5c08613

Simpson, E. H. (1949). Measurement of Diversity. Nature, 163(4148), Article 4148. 10.1038/163688a0

Smith, D. (2026). rbiom: Integrated Analysis and Visualization of Microbiome Data. https://github.com/cmmr/rbiom

Tak, P. P., & Firestein, G. S. (2001). NF-κB: A key role in inflammatory diseases. The Journal of Clinical Investigation, 107(1), 7–11. 10.1172/JCI11830

Takada, T., Kurakawa, T., Tsuji, H., & Nomoto, K. (2013). Fusicatenibacter saccharivorans gen. Nov., sp. Nov., isolated from human faeces. International Journal of Systematic and Evolutionary Microbiology, 63(Pt_10), 3691–3696. 10.1099/ijs.0.045823-0

Takeshita, K., Mizuno, S., Mikami, Y., Sujino, T., Saigusa, K., Matsuoka, K., Naganuma, M., Sato, T., Takada, T., Tsuji, H., Kushiro, A., Nomoto, K., & Kanai, T. (2016). A Single Species of Clostridium Subcluster XIVa Decreased in Ulcerative Colitis Patients. Inflammatory Bowel Diseases, 22(12), 2802–2810. 10.1097/MIB.0000000000000972

Tang, H., Huang, W., & Yao, Y.-F. (2023). The metabolites of lactic acid bacteria: Classification, biosynthesis and modulation of gut microbiota. Microbial Cell, 10(3), 49–62. 10.15698/mic2023.03.792

Tang, Q., Jin, G., Wang, G., Liu, T., Liu, X., Wang, B., & Cao, H. (2020). Current Sampling Methods for Gut Microbiota: A Call for More Precise Devices. Frontiers in Cellular and Infection Microbiology, 10. 10.3389/fcimb.2020.00151

Turpin, W., Espin-Garcia, O., Xu, W., Silverberg, M. S., Kevans, D., Smith, M. I., Guttman, D. S., Griffiths, A., Panaccione, R., Otley, A., Xu, L., Shestopaloff, K., Moreno-Hagelsieb, G., Paterson, A. D., & Croitoru, K. (2016). Association of host genome with intestinal microbial composition in a large healthy cohort. Nature Genetics, 48(11), Article 11. 10.1038/ng.3693

Ulluwishewa, D., Anderson, R. C., McNabb, W. C., Moughan, P. J., Wells, J. M., & Roy, N. C. (2011). Regulation of Tight Junction Permeability by Intestinal Bacteria and Dietary Components1,2. The Journal of Nutrition, 141(5), 769–776. 10.3945/jn.110.135657

Villmones, H. C., Haug, E. S., Ulvestad, E., Grude, N., Stenstad, T., Halland, A., & Kommedal, Ø. (2018). Species Level Description of the Human Ileal Bacterial Microbiota. Scientific Reports, 8(1), 4736. 10.1038/s41598-018-23198-5

Walker, S. P., Barrett, M., Hogan, G., Flores Bueso, Y., Claesson, M. J., & Tangney, M. (2020). Non-specific amplification of human DNA is a major challenge for 16S rRNA gene sequence analysis. Scientific Reports, 10, 16356. 10.1038/s41598-020-73403-7

Wang, J., Thingholm, L. B., Skiecevičienė, J., Rausch, P., Kummen, M., Hov, J. R., Degenhardt, F., Heinsen, F.-A., Rühlemann, M. C., Szymczak, S., Holm, K., Esko, T., Sun, J., Pricop-Jeckstadt, M., Al-Dury, S., Bohov, P., Bethune, J., Sommer, F., Ellinghaus, D., … Franke, A. (2016). Genome-wide association analysis identifies variation in vitamin D receptor and other host factors influencing the gut microbiota. Nature Genetics, 48(11), Article 11. 10.1038/ng.3695

Wang, M., Ahrné, S., Jeppsson, B., & Molin, G. (2005). Comparison of bacterial diversity along the human intestinal tract by direct cloning and sequencing of 16S rRNA genes. FEMS Microbiology Ecology, 54(2), 219–231. 10.1016/j.femsec.2005.03.012

Wang, M., Ma, W., Wang, C., & Li, D. (2024). Lactococcus G423 improve growth performance and lipid metabolism of broilers through modulating the gut microbiota and metabolites. Frontiers in Microbiology, 15, 1381756. 10.3389/fmicb.2024.1381756

Wang, Y., & LêCao, K.-A. (2020). Managing batch effects in microbiome data. Briefings in Bioinformatics, 21(6), 1954–1970. 10.1093/bib/bbz105

Witten, D. M., Tibshirani, R., & Hastie, T. (2009). A penalized matrix decomposition, with applications to sparse principal components and canonical correlation analysis. Biostatistics, 10(3), 515–534. 10.1093/biostatistics/kxp008

Witten, D. M., & Tibshirani, R. J. (2009). Extensions of Sparse Canonical Correlation Analysis with Applications to Genomic Data. Statistical Applications in Genetics and Molecular Biology, 8(1), 28. 10.2202/1544-6115.1470

Wu, T., Hu, E., Xu, S., Chen, M., Guo, P., Dai, Z., Feng, T., Zhou, L., Tang, W., Zhan, L., Fu, X., Liu, S., Bo, X., & Yu, G. (2021). clusterProfiler 4.0: A universal enrichment tool for interpreting omics data. The Innovation, 2(3), 100141. 10.1016/j.xinn.2021.100141

Yamamoto, R., Chung, R., Vazquez, J. M., Sheng, H., Steinberg, P. L., Ioannidis, N. M., & Sudmant, P. H. (2022). Tissue-specific impacts of aging and genetics on gene expression patterns in humans. Nature Communications, 13(1), 5803. 10.1038/s41467-022-33509-0

Yang, K., Li, G., Li, Q., Wang, W., Zhao, X., Shao, N., Qiu, H., Liu, J., Xu, L., & Zhao, J. (2025). Distribution of gut microbiota across intestinal segments and their impact on human physiological and pathological processes. Cell & Bioscience, 15(1), 47. 10.1186/s13578-025-01385-y

Yersin, S., & Vonaesch, P. (2024). Small intestinal microbiota: From taxonomic composition to metabolism. Trends in Microbiology, 32(10), 970–983. 10.1016/j.tim.2024.02.013

Yoda, M., Nomura, N., Yoda, S., Kagotani, M., Murakami, A., Namai, F., Fujii, T., Tochio, T., Sato, T., & Shimosato, T. (2026). Genetically Modified Lactococcus lactis Hypersecreting IL-1Ra Improves Glucose Metabolism and Modulates the Gut Microbiota in an Obese Mouse Model. Journal of Diabetes Research, 2026, 6006491. 10.1155/jdr/6006491

Yu, G., Wang, L.-G., Han, Y., & He, Q.-Y. (2012). clusterProfiler: An R Package for Comparing Biological Themes Among Gene Clusters. OMICS: A Journal of Integrative Biology, 16(5), 284–287. 10.1089/omi.2011.0118

Yue, X., Wen, S., Long-kun, D., Man, Y., Chang, S., Min, Z., Shuang-yu, L., Xin, Q., Jie, M., & Liang, W. (2022). Three important short-chain fatty acids (SCFAs) attenuate the inflammatory response induced by 5-FU and maintain the integrity of intestinal mucosal tight junction. BMC Immunology, 23(1), 19. 10.1186/s12865-022-00495-3

Zaharia, M., Bolosky, W. J., Curtis, K., Fox, A., Patterson, D., Shenker, S., Stoica, I., Karp, R. M., & Sittler, T. (2011). *Faster and More Accurate Sequence Alignment with SNAP* (arXiv:1111.5572). arXiv. 10.48550/arXiv.1111.5572

Zheng, D., Liwinski, T., & Elinav, E. (2020). Interaction between microbiota and immunity in health and disease. Cell Research, 30(6), 492–506. 10.1038/s41422-020-0332-7

Zhou, W., Sailani, M. R., Contrepois, K., Zhou, Y., Ahadi, S., Leopold, S. R., Zhang, M. J., Rao, V., Avina, M., Mishra, T., Johnson, J., Lee-McMullen, B., Chen, S., Metwally, A. A., Tran, T. D. B., Nguyen, H., Zhou, X., Albright, B., Hong, B.-Y., … Snyder, M. (2019). Longitudinal multi-omics of host–microbe dynamics in prediabetes. Nature, 569(7758), Article 7758. 10.1038/s41586-019-1236-x

