## Supplementary figures and images for "Integrative host transcriptomic and mucosal microbiome profiling reveals region-specific host-microbiome associations across the human intestine"

### Supplemental Figure 1

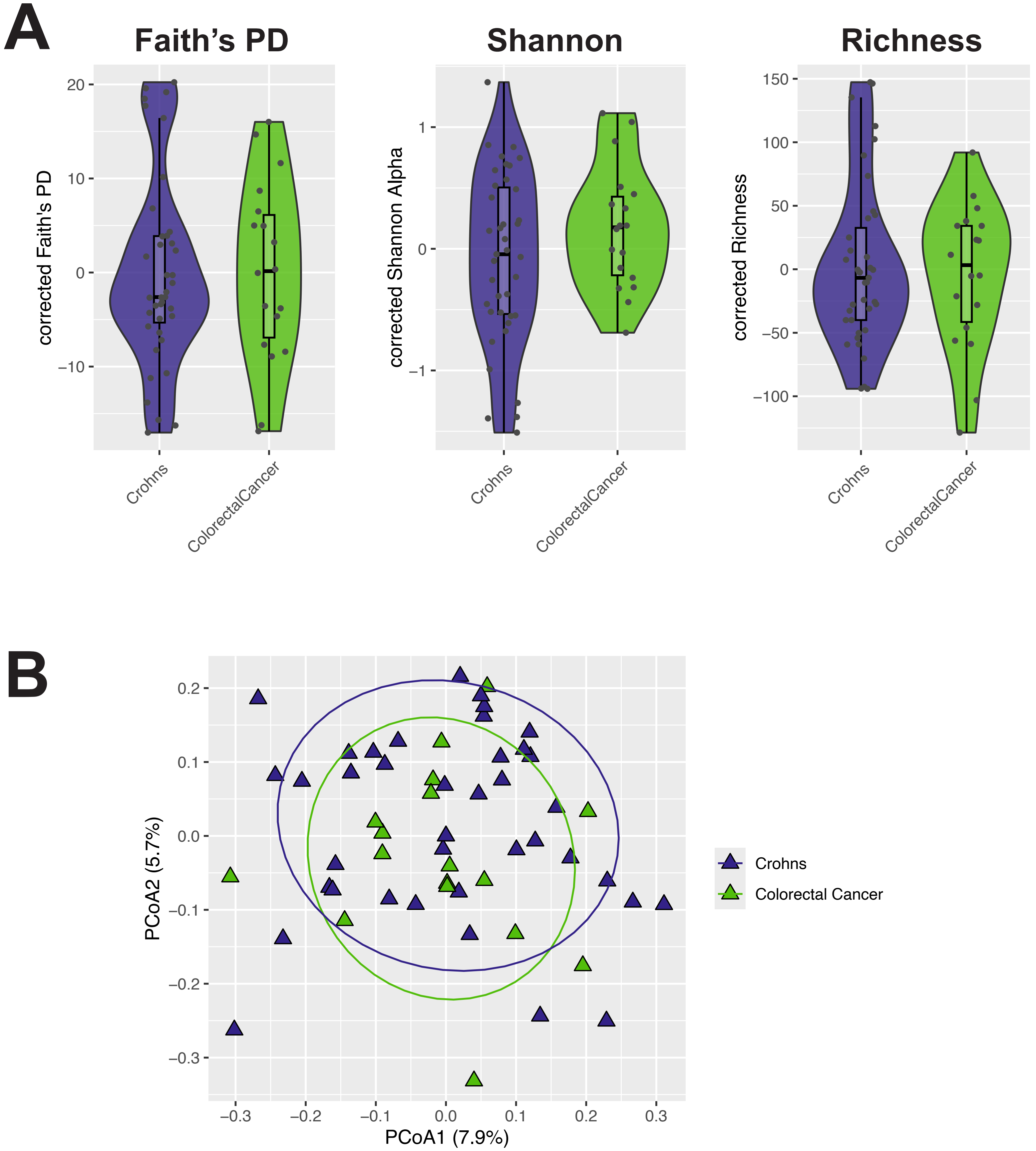

### Supplemental Figure 2

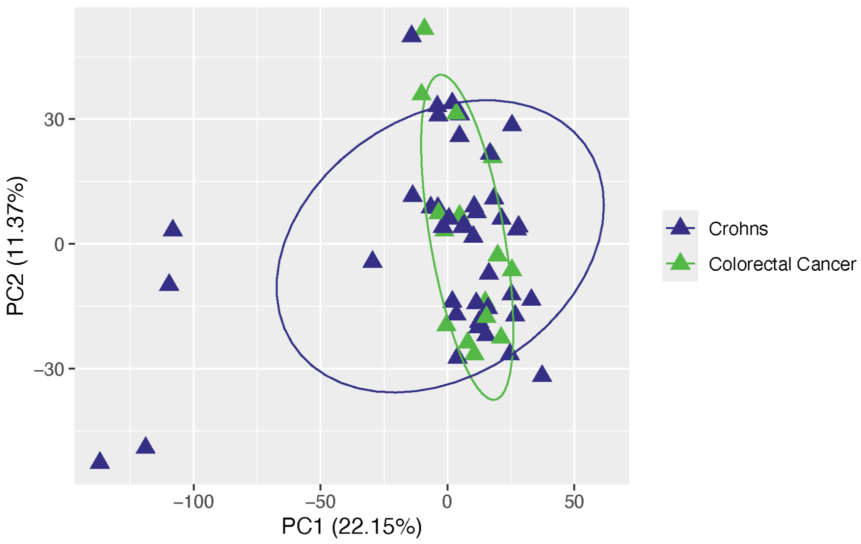

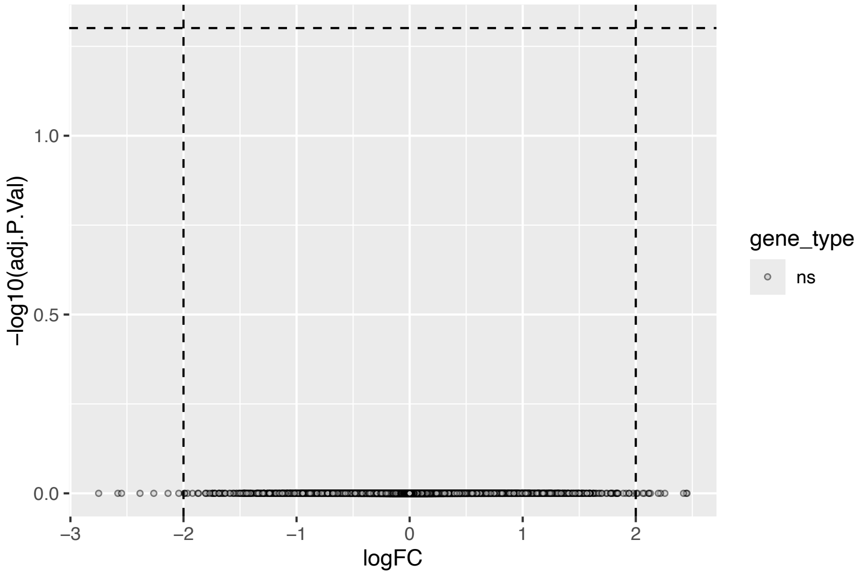

### Supplemental Figure 3

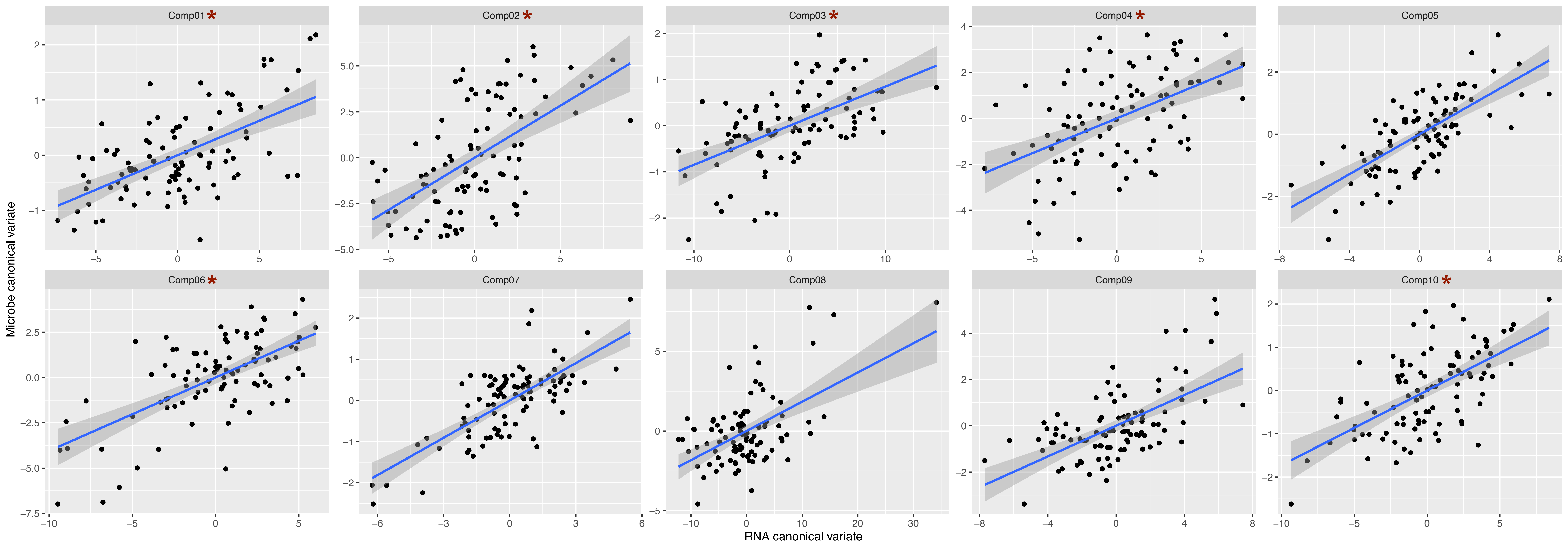

### Supplemental Figure 4

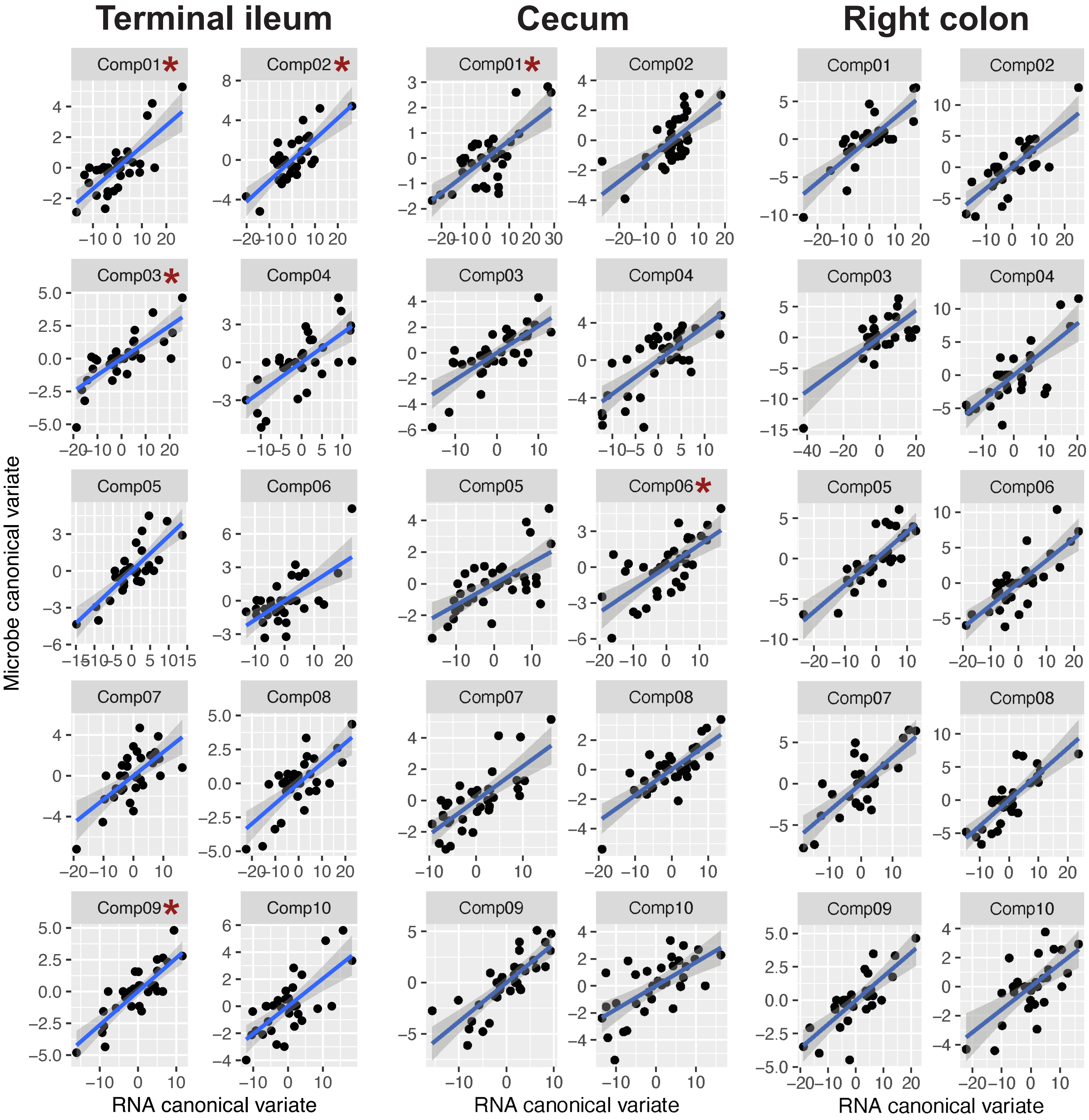

### Supplemental Figure 5

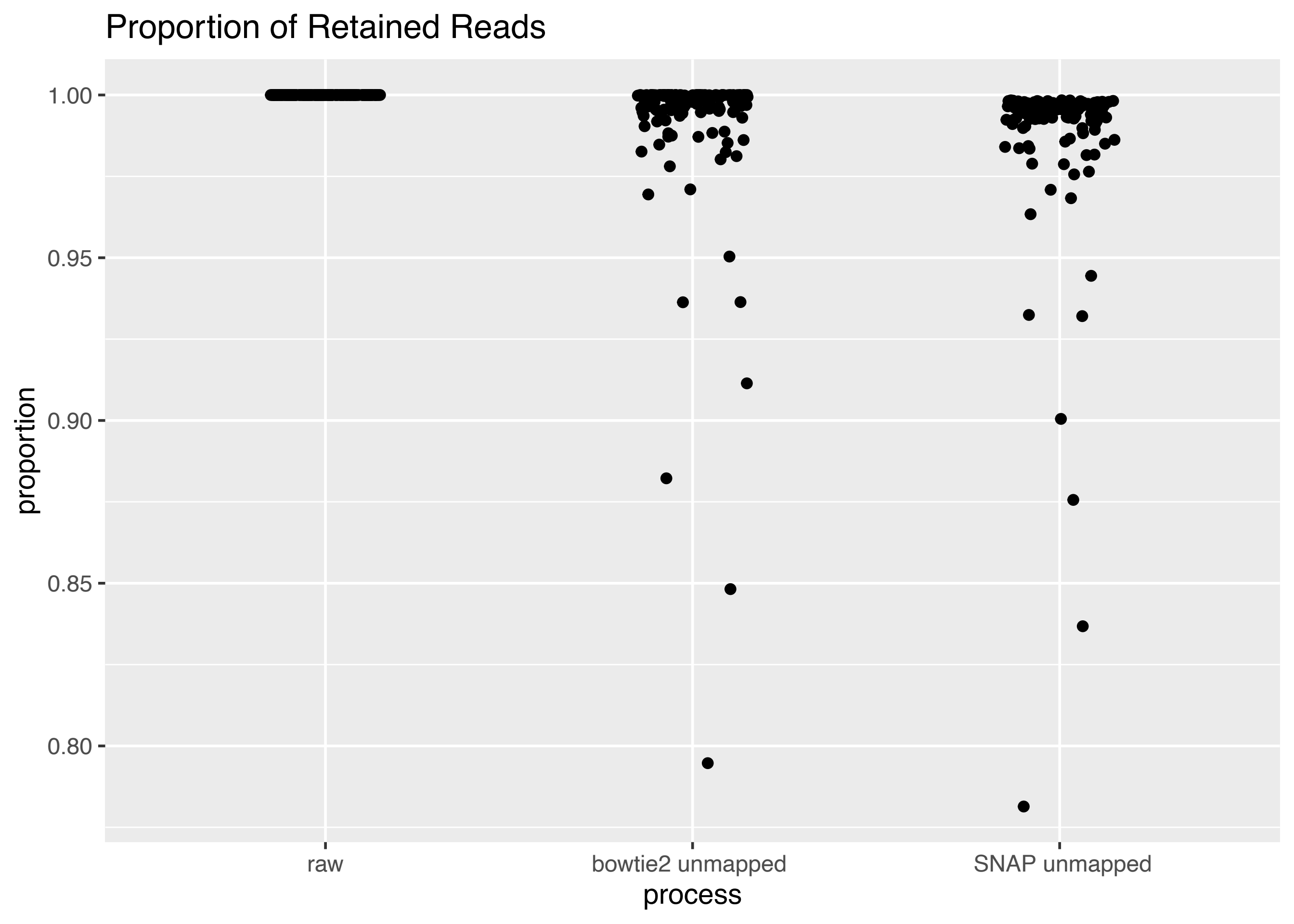

### Supplemental Figure 6

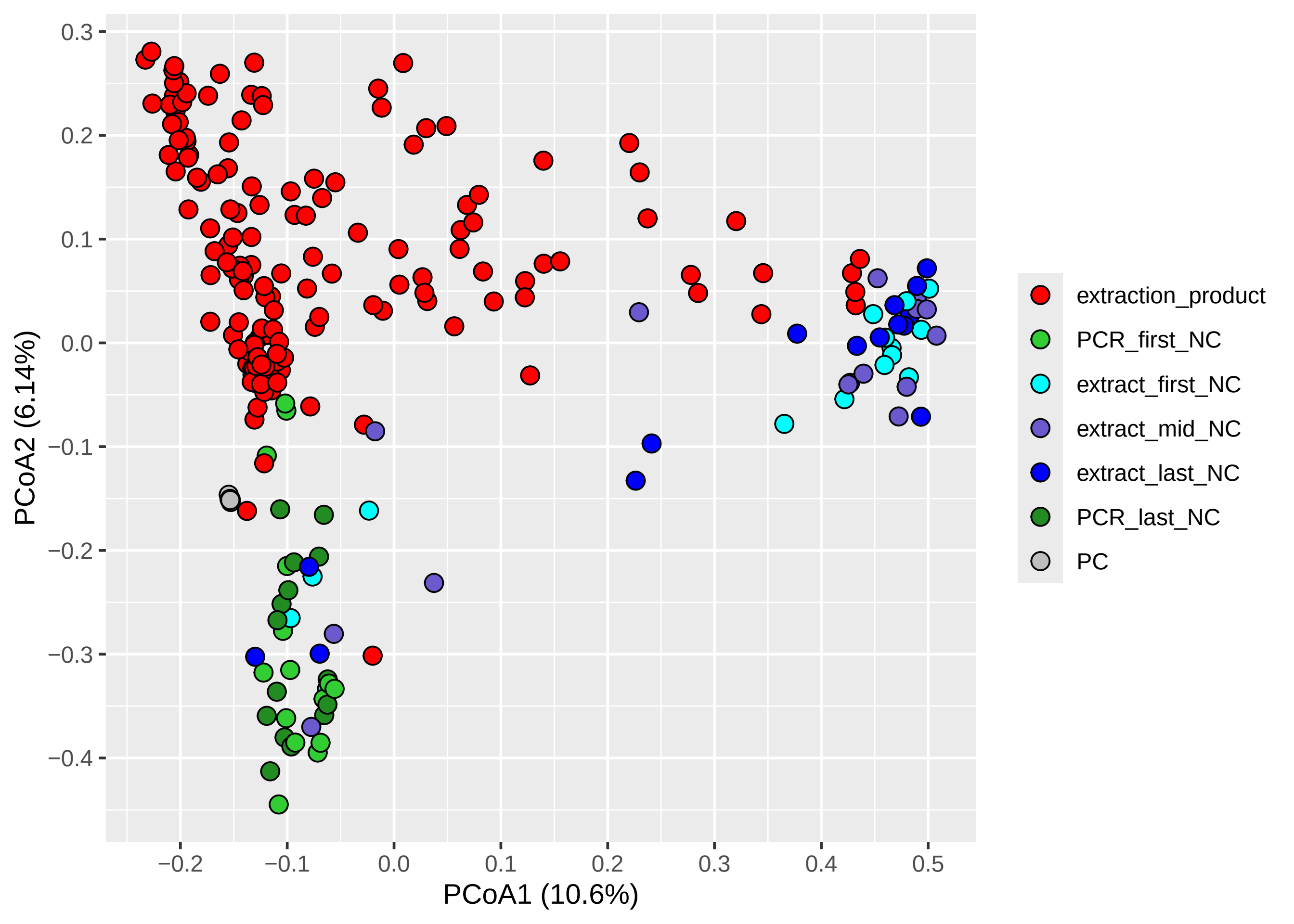

### Supplemental Figure 7

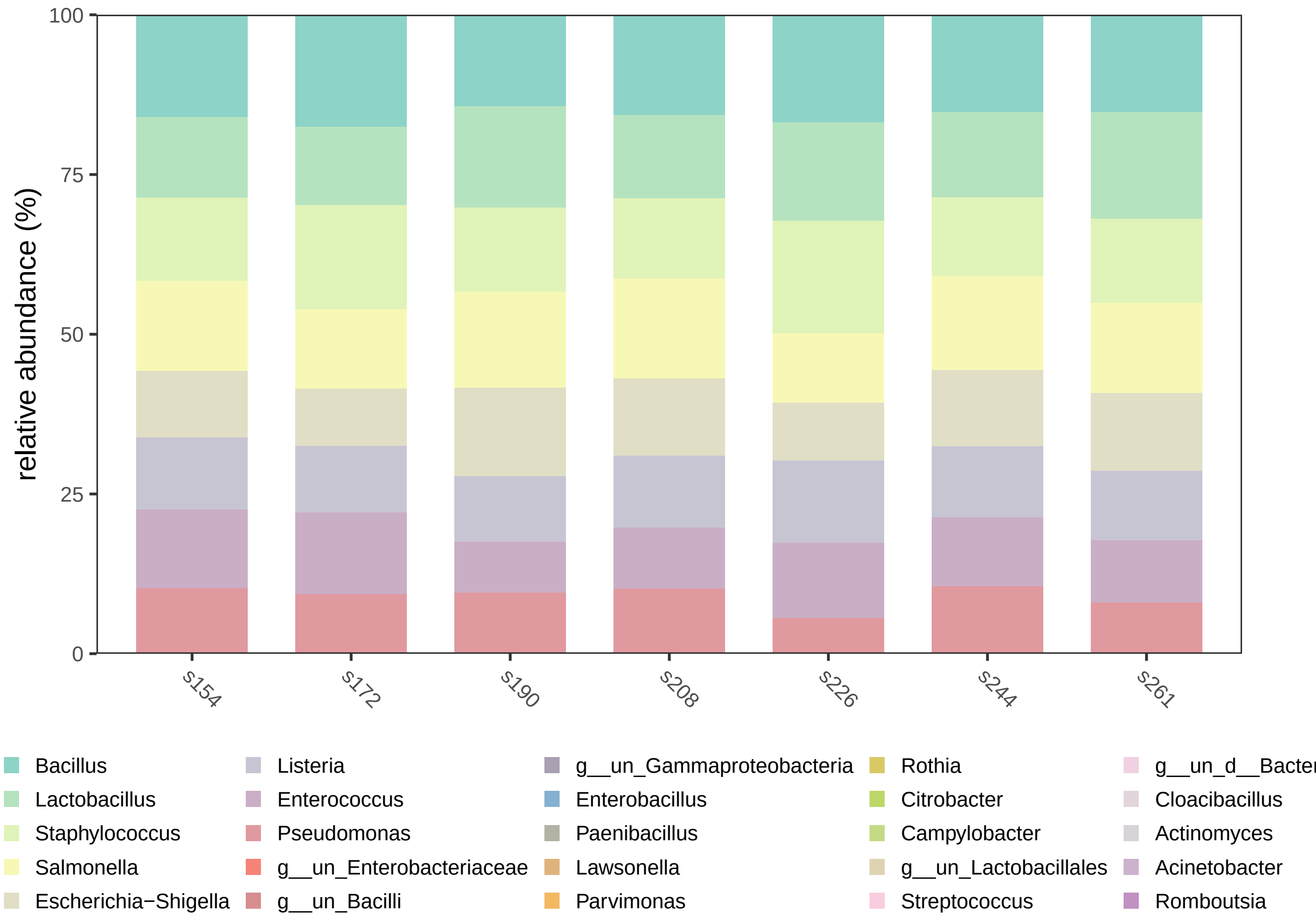

### Supplemental Figure 8

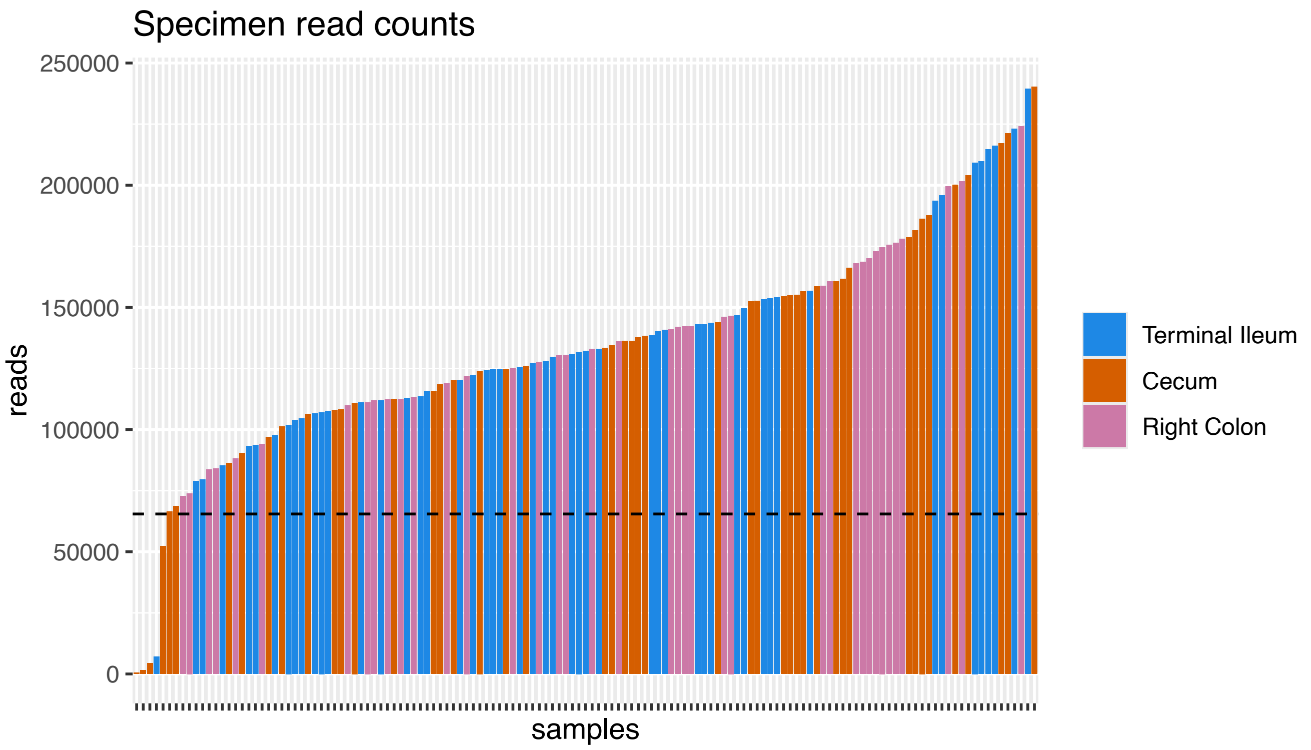

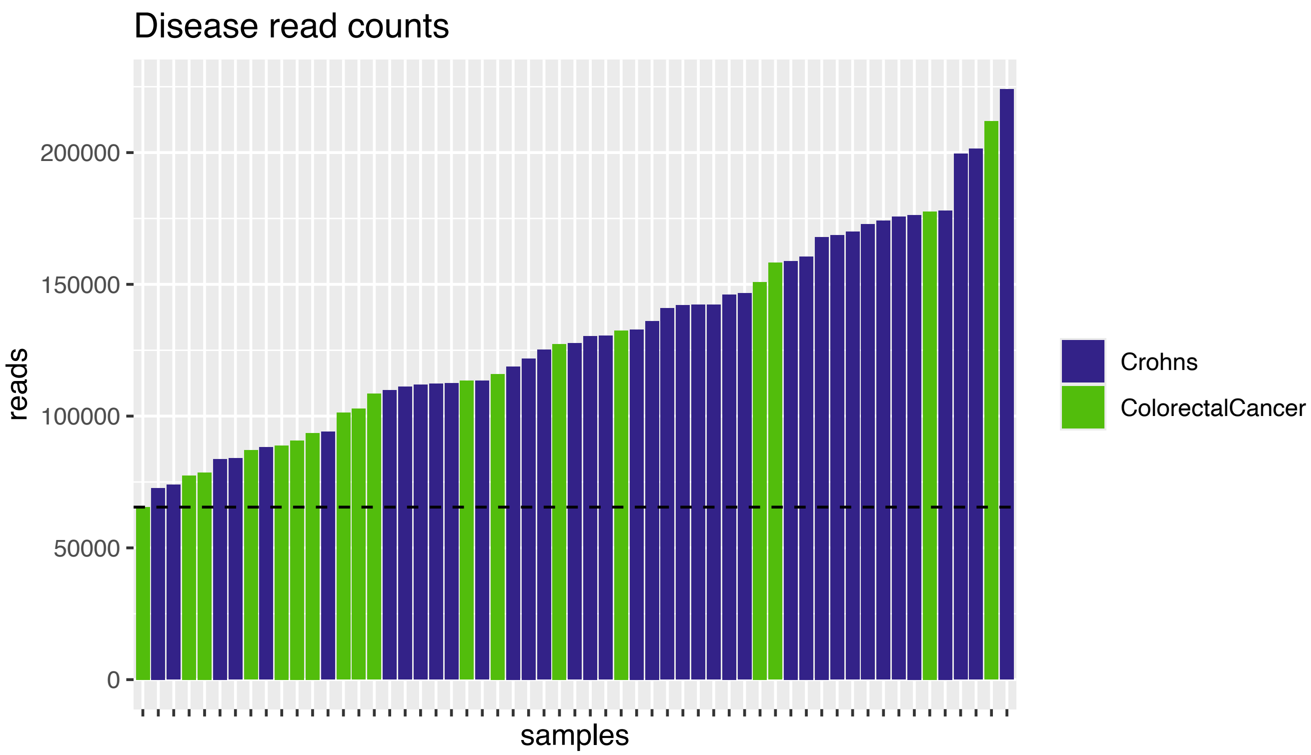
